# Somatic embryogenesis of grapevine (*Vitis vinifera*) expresses a transcriptomic hourglass

**DOI:** 10.1101/2024.04.08.588272

**Authors:** Sara Koska, Dunja Leljak-Levanić, Nenad Malenica, Kian Bigović Villi, Momir Futo, Nina Čorak, Mateja Jagić, Anja Tušar, Niko Kasalo, Mirjana Domazet-Lošo, Kristian Vlahoviček, Tomislav Domazet-Lošo

## Abstract

At the molecular level, multicellular eukaryotic lineages and bacterial biofilms show predictable evolutionary footprints in their development. For instance, the zygotic embryogenesis of *Arabidopsis*, which is initiated by gamete fusion, shows hourglass-shaped ontogeny-phylogeny correlations at the transcriptome level. However, many plants are capable of yielding a fully viable next generation by somatic embryogenesis — a comparable developmental process that usually starts by the embryogenic induction of a diploid somatic cell. This leads to the question: is the hourglass-shaped ontogeny-phylogeny correlation preserved in somatic embryogenesis? To explore the correspondence between ontogeny and phylogeny in this alternative developmental route in plants, we developed a new and highly efficient model of somatic embryogenesis in grapevine (*Vitis vinifera*) and sequenced its developmental transcriptomes. By combining the evolutionary properties of grapevine genes with their expression values, which were recovered from early induction until the formation of juvenile plants, we found a strongly supported hourglass-shaped developmental trajectory. However, in contrast to zygotic embryogenesis in *Arabidopsis* where the torpedo stage was evolutionary the most inert, we found that in the somatic embryogenesis of grapevine the heart stage expressed evolutionary the oldest and the most conserved transcriptome. This is a surprising finding because it suggests a better evolutionary system-level analogy between animal development and plant somatic embryogenesis than zygotic embryogenesis. We conclude that macroevolutionary logic is deeply hardwired in plant ontogeny and that somatic embryogenesis is likely a primordial embryogenic program in plants.

## Introduction

The ontogenies of multicellular eukaryotes are commonly marked by macroevolutionary imprints at the molecular level (Domazet-Lošo and Tautz 2010b; Kalinka et al. 2010; Irie and Kuratani 2011; Quint et al. 2012; Cheng et al. 2015; Drost et al. 2015; Drost et al. 2016). Curiously, we recently found that similar regularities are also present in the development of bacterial biofilms; a process that mimics embryogenesis of multicellular eukaryotes (Futo et al. 2021). However, an hourglass-shaped correlation between ontogeny and phylogeny is a unique feature of multicellular eukaryotes. This pattern was first discovered in various animals by several independent approaches that compared the evolutionary conservation of genes and the ontogenetic timing of their expression (Domazet-Lošo and Tautz 2010b; Kalinka et al. 2010; Irie and Kuratani 2011). Subsequently, the hourglass-shaped ontogeny-phylogeny correspondence was also discovered in the transcriptomes of a plant species *Arabidopsis thaliana* (Quint et al. 2012; Drost et al. 2015; Drost et al. 2016). This was a largely unexpected finding because embryogenesis in plants, in contrast to animals, does not hint at the existence of such correlations at the morphological level (Drost et al. 2016, Drost et al. 2017). However, so far correlation between ontogeny and phylogeny in plants has been only explored in the zygotic embryogenesis of *A. thaliana* (Quint et al. 2012; Drost et al. 2015; Drost et al. 2016), leaving the existence of such correlations in other plant species or alternative developmental routes completely uncertain.

The organismal development of both animals and plants usually starts with the zygote formation and unfolds through the process of embryogenesis. In flowering plants, zygotic embryogenesis (ZE) involves double fertilization following the simultaneous formation of the embryo and the endosperm within a developing seed after a set of morphological, cellular, and molecular changes (Radoeva and Weijers 2014). However, in contrast to animals that mainly have sexual ontogeny, life cycle in plants includes another level of complexity in the form of somatic embryogenesis (SE). During this process plant embryos develop from cells other than the fertilized eggs (Dodeman et al. 1997; Horstman et al. 2017). This is an alternative road to embryo-mediated plant formation, which typically includes reprogramming of somatic cells towards the embryogenic pathway (Williams and Maheswaran 1986; Brukhin and Morozova 2011; Smertenko and Bozhkov 2014). This process, in contrast to zygotic embryogenesis, generates embryos which are genetically identical to the mother plant (Salaün et al. 2021). The best-known example of naturally occurring somatic embryogenesis is the genus *Kalanchoe*, commonly called the mother of thousands, in which somatic embryos form spontaneously from diploid somatic cells on leaves edges (Dodeman et al. 1997; Garcês and Sinha 2009).

In many plant species, somatic embryogenesis can be artificially induced in different cell types after exposing the cells to various SE-promoting conditions (Dodeman et al. 1997; Smertenko and Bozhkov 2014; Horstman et al. 2017). Akin to a zygote, a dedifferentiated somatic cell that initiates somatic embryogenesis usually shows cell polarity (Smertenko and Bozhkov 2014; Mendez-Hernandez et al. 2019). The subsequent development of somatic embryos, at least in *Arabidopsis* and other dicots, roughly follows morphological transitions characteristic for zygotic embryogenesis; i.e., globular, heart, torpedo, and cotyledonary stages (Kurczyńska et al. 2007). At the molecular level, some key developmental regulators, such as BBM and SERK1 (Smertenko and Bozhkov 2014), are shown to be used both in somatic and zygotic embryogenesis. Actually, some transcription factors like FUS3 and AGL15, which play a central role in zygotic embryogenesis, are also involved in the ectopic embryo initiation of somatic embryogenesis (Radoeva and Weijers 2014).

On the other hand, the currently available comparative transcriptome analyses of somatic and zygotic embryos, although limited to only a few developmental stages, reveal substantial transcriptional differences between these two developmental routes for a huge number of genes (Jin et al. 2014; Maximova et al. 2014, Hofmann et al. 2019). These large disparities between ZE and SE transcriptomes are perhaps not surprising given that many striking differences between zygotic and somatic embryogenesis exist at the morphological and functional levels. For example, zygotic embryo development occurs inside the seed, where intensive communication between the embryo and surrounding endosperm occurs (Berger et al. 2006; Zhou et al. 2013). In somatic embryogenesis, these interactions are missing because double fertilization, and consequently endosperm formation, are absent (Leljak-Levanić et al. 2015; Matilla 2019). Similarly, zygotic embryos go through the phase of metabolic quiescence and desiccation, which is a part of seed maturation known as seed dormancy (Braybrook and Harada 2008). However, somatic embryos undergo the full developmental trajectory in the absence of these processes.

Furthermore, somatic embryos are larger in size and their cells are metabolically more active, contain bigger vacuoles, more lipid droplets, and mitochondria in comparison to zygotic embryos (Jin et al. 2014). Interestingly, zygotic embryos with impaired endosperm development in rice and *Arabidopsis* also show increased total embryo size (Hong et al. 1996; Guo et al. 2010; Ando et al. 2023). Unlike zygotic embryogenesis, somatic embryogenesis is triggered by a wider range of factors including stress factors like the wounding of explant tissues or the high concentrations of plant growth regulators such as auxin (Jin et al. 2014; Horstman et al. 2017; Mendez-Hernandez et al. 2019). All of this implies that somatic embryogenesis, despite similar final developmental outcomes compared to zygotic embryogenesis, is a unique developmental route in seed plants.

For instance, plant embryos derived from fertilization are very difficult to isolate in their earliest stages due to their small size and enclosure within seeds (Dodeman et al. 1997; Xiang et al. 2011; Venglat et al. 2013). Although there are many studies encompassing transcriptomes of zygotic embryogenesis from the pre-globular stages onward (Chen et al. 2014; Jin et al. 2014; Drost et al. 2016; Itoh et al. 2016; Silva et al. 2016; Hofmann et al. 2019), very little is known about the molecular aspects in the very first steps of zygotic embryo development following fertilization (Xiang et al. 2011; Venglat et al. 2013). In this context, somatic embryogenesis has a great advantage because somatic embryos are accessible at any stage of their development, which makes somatic embryogenesis a favorable model of plant embryogenesis (Dodeman et al 1997; Fehér et al. 2003). Another advantage of somatic embryogenesis is that it allows an easy clonal propagation which is helpful in situations where efficient production of large numbers of genetically identical plants is required (Smertenko et al. 2014; Salaün et al. 2021).

Several studies explored transcriptomes of somatic embryos in different plant species (Lin et al. 2009; Jin et al. 2014; Maximova et al. 2014; Wickramasuriya and Dunwell 2015; Magnani et al. 2017; Kang et al. 2021; Ci et al. 2022). Unfortunately, these studies focus on a single or a few stages of somatic embryogenesis, thus covering only a fraction of this developmental process (Jin et al. 2014; Maximova et al. 2014; Wickramasuriya and Dunwell 2015; Magnani et al. 2017; Kang et al. 2021). In addition, RNA samples in some of these studies are derived from a mixture of different SE developmental stages (Lin et al. 2009; Wickramasuriya and Dunwell 2015; Magnani et al. 2017; Kang et al. 2021; Ci et al. 2022), which precludes the recovery of time-resolved transcriptional trajectories. These limitations of currently available SE datasets make them unsuitable for studying phylogeny-ontogeny correlations, given that this type of analysis requires relatively dense sampling of individual stages along the full developmental process (Futo et al. 2022).

To overcome these limitations and to explore phylogeny-ontogeny correlations along the full SE developmental process, we established a highly efficient protocol for the direct induction of somatic embryogenesis in grapevine (*Vitis vinifera* L.) ‘Malvasia Istriana’, a perennial woody dicotyledon species and a cultivar with an international reputation. The developmental stages of our *V. vinifera* somatic embryogenesis morphologically roughly resemble the stages of normal zygotic embryogenesis, and the resulting embryos possess a high potential for immediate plantlet regeneration. We used this novel system to sample 12 morphologically distinct developmental stages, covering the full ontogeny of *V. vinifera* somatic embryogenesis from early induction until the formation of a juvenile plant, and sequenced their transcriptomes by RNAseq. We matched the obtained expression values to gene-related evolutionary and functional information to explore the correspondence between ontogeny and phylogeny along the SE developmental trajectory. To achieve this, we applied phylostratigraphic and phylotranscriptomic methods which are very powerful in extracting macroevolutionary information from genomic and developmental data (Domazet-Lošo et al. 2007, Domazet-Lošo and Tautz 2010a, Domazet-Lošo and Tautz 2010b, Quint et al. 2012, Domazet-Lošo et al. 2017, Drost et al. 2018, Futo et al. 2021, Domazet-Lošo et al. 2024).

Here we show a strongly supported hourglass-shaped developmental trajectory in *V. vinifera* somatic embryogenesis. Moreover, the recovered somatic embryogenesis patterns better align with theoretical expectations than previously found profiles in zygotic embryogenesis. This suggests that somatic embryogenesis is a primordial process tightly linked to the evolutionary origin of development in plants.

## Results

### Global expression profiles along SE

To evolutionary assess developmental transcriptomes of somatic embryogenesis, we first developed a new and highly efficient SE induction system in grapevine (*Vitis vinifera* L.) ‘Malvasia Istriana’, which is characterized by a high embryogenic potential, developmental synchronization between embryos after induction, and the absence of fusion between embryos at their interfaces (see Methods). These properties of our SE system allowed us to relatively easily select and isolate individual embryos at different developmental stages in sufficient amounts for downstream RNAseq analysis (Figure 1a). Embryogenesis was induced from immature anthers which are the most reactive explants for SE in different grapevine cultivars (Cadavid-Labrada et al. 2008, Dhekney et al. 2009, Malenica et al. 2020). To cover the full embryogenesis as well as post-embryonic development, we used three cultivation media and different lighting conditions that simulate dormancy and germination (Figure 1a). This allowed us to gather a collection of 12 SE developmental stages covering induction, embryonic, and postembryonic development, including plantlet formation (Figure 1a). To our knowledge, this is the most complete SE sample collection generated so far.

To get an overview of expression dynamics along SE in *V. vinifera,* we recovered the transcriptomes of these 12 SE stages by RNAseq (Figure 1a). If we consider all sequenced developmental stages together, we detected expression for 29,839 (99.56%) annotated *V. vinifera* genes (Supplementary Data 1). These high numbers showed that essentially all genes were transcribed at some point along the developmental trajectory of somatic embryogenesis. In turn, this reveals that somatic embryogenesis cannot be considered some simplified derivative of zygotic embryogenesis, but rather a full-fledged developmental process that utilizes essentially all available protein-coding genetic information.

We further tested expression dynamics along the whole SE ontogeny which revealed that 25,098 (84.1 %) genes were differentially expressed (Supplementary Data 2). Among these 12,893 (43.2 %) had expression change above two-fold. To get a more detailed overview of expression dynamics we also compared transcriptomes in a pairwise manner between successive stages (Supplementary Data 2 and Supplementary Figure 1). This pairwise analysis revealed that on average 18 % of genes (12 %, fold change > 2) changed expression in transition between two successive stages. The most dramatic change occurs at the transition from C1 to C2 stage where 37 % (25 %, fold change > 2) genes change expression (Supplementary Data 2 and Supplementary Figure 1). These values, together with clustering analysis (Supplementary Data 3 and 4), revealed that somatic embryogenesis in *V. vinifera* is a highly regulated process underpinned by substantial changes at the transcriptome level.

To get a global overview of the similarities and differences between the expressed transcriptomes of different developmental stages, we calculated pair-wise expression correlations, which revealed that the SE developmental trajectory could be divided into five expression phases (Figure 1a, b). The early expression phase includes the early induction (EI), pre-globular (PG), and globular (G1, G2) development stages (Figure 1a, b). The mid expression phase covers the heart (H) and early torpedo (T1) developmental stages. This mid expression phase is followed by the late torpedo (T2) and early cotyledonary (C1) developmental stages which have rather unique transcriptomes that show some discontinuous similarity to the mid and late expression phase (Figure 1b). Finally, the late expression phase comprises late cotyledonary (C2), the formation of seedling (S), seedling with epicotyl (EP) and juvenile plant (JP) developmental stages (Figure 1a, b).

**Figure 1.**
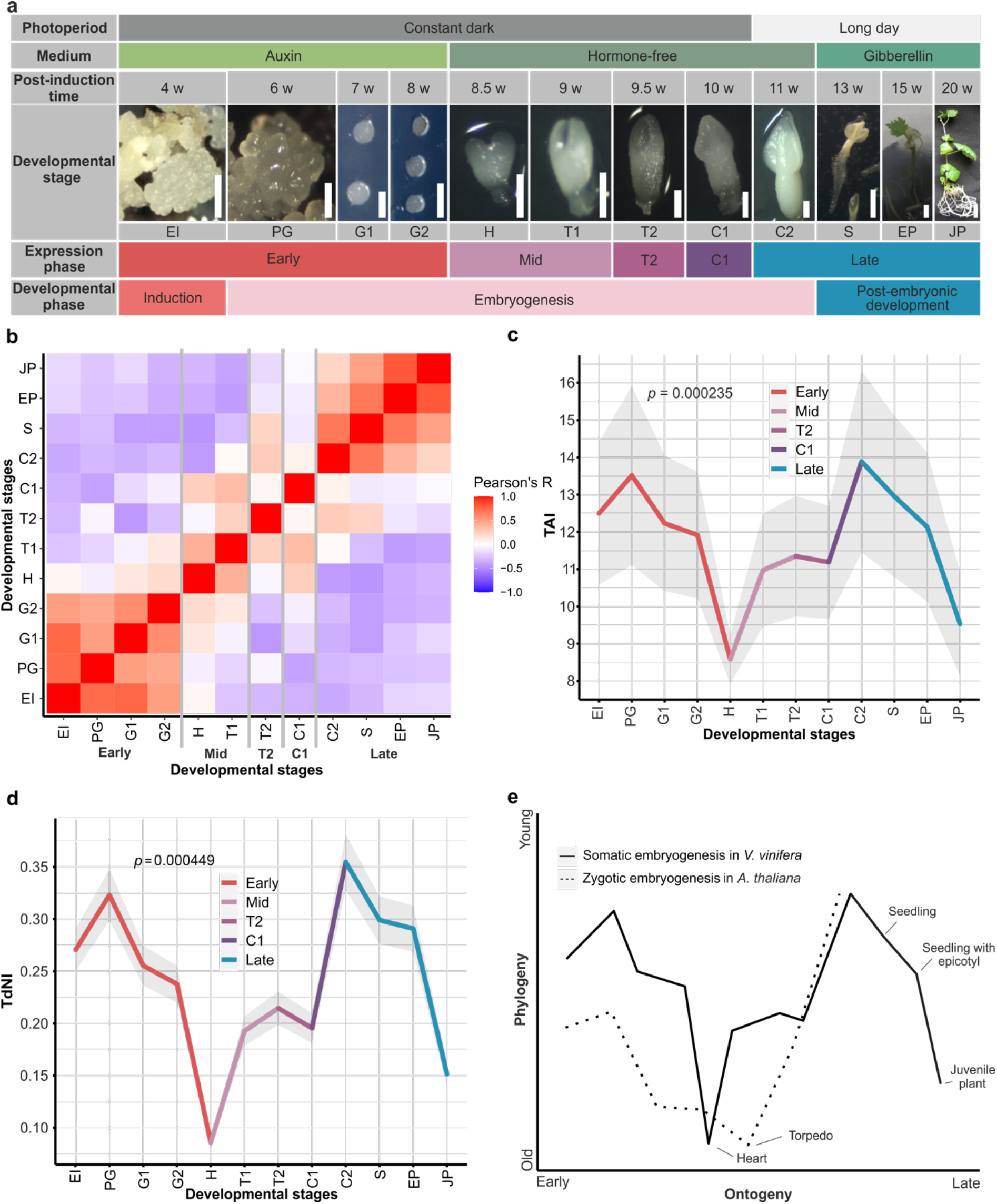
Somatic embryogenesis in *V. vinifera* is a stage-organized process that exhibits an hourglass-shaped phylogeny-ontogeny correlation. **a**) The sampled developmental stages of somatic embryogenesis in *V. vinifera*: early induction (EI), preglobular stage (PG), globular stage 1 (G1), globular stage 2 (G2), heart stage (H), torpedo stage 1 (T1), torpedo stage 2 (T2), cotyledonary stage 1 (C1), cotyledonary stage 2 (C2), seedling (S), seedling with epicotyl (EP) and juvenile plant (JP). Size bars: 0.5 mm (EI – C2), 2 mm (S), 3 mm (EP), 1 cm (JP). The somatic embryogenesis stages were determined following previously described morphological criteria (Malenica et al. 2020). For every sampled developmental stage, we showed corresponding hormones that were present in media as well as photoperiod at which developing plants were cultivated. “Long day” marks photoperiod of 18h light and 6h dark. For an easy reference, we also depicted post-induction time in weeks (w), global developmental phases and expression phases derived from our correlation analysis. **b)** Pearson’s correlation coefficients between somatic embryogenesis developmental stages in all-against-all comparison. Early (EI – G2), mid (H – T1), T2, C1 and late (C2 – JP) expression stages are marked. **c)** The transcriptome age index (TAI) of somatic embryogenesis shows an hourglass pattern. The heart stage of the mid developmental period expresses the evolutionary oldest transcriptome, while earlier and later stages express evolutionary younger ones. We tested the significance of the TAI pattern using the flat line test, while the grey shaded area represents ± one standard deviation estimated using permutation analysis (see Methods). **d)** The transcriptome non-synonymous divergence index (TdNI) of somatic embryogenesis shows an hourglass pattern. The heart stage of the mid developmental period expresses the most conserved genes at non-synonymous divergence sites, while earlier and later stages express more diverged genes. Non-synonymous divergence rates were estimated in *V. vinifera – V. arizonica* pairwise comparisons (see Methods). We tested the significance of the TdNI pattern using the flat line test, while the grey shaded area represents ± one standard deviation estimated using permutation analysis (see Methods). The corresponding transcriptome synonymous divergence index (TdSI) and transcriptome codon bias index (TCBI) profiles are shown in Supplementary Figure 2e. A schematic comparison between the TAI profile of *V. vinifera* somatic embryogenesis that we recovered in this study and the TAI profile of *A. thaliana* zygotic embryogenesis reported previously (Quint et al. 2012). To make the hourglass patterns visually comparable between these studies, the TAI values were normalized to a range between 0 and 1 (see Methods).

To get further insights into expression dynamics along the SE developmental trajectory, we performed principal component analysis (PCA) which revealed a time-resolved profile that follows the developmental progression of somatic embryogenesis and shows its punctuated organization (Figure 2). The general organization of this PCA pattern in SE is similar to those previously recovered in bacterial biofilm development (Futo et al. 2021). This suggests that these developmental processes, although analogous, are governed by the common basic principles. Similar to bacterial biofilm development (Futo et al. 2021), biological replicates per developmental stage generally clustered together (Figure 2). The only stage that showed increased distortion in expression between replicates is cotyledonary stage 1 (C1). This pattern in C1 could reflect a burst of expression changes, which potentially could be resolved in future studies by even finer temporal sampling around this specific period. Alternatively, this could point to an increased sensitivity of this particular stage to the slight changes in environmental cues. We previously observed similar patterns during biofilm growth in developmental stages which were impacted by substantial environmental stress caused by starvation (Futo et al. 2021). The fact that C1 stage is the latest stage kept in constant dark (Figure 1a) — which causes a tradeoff between the lack of photosynthesis and developmental growth — suggests that higher variability in transcriptomes between biological replicates in C1 stage likely reflects the effect of starvation (Figure 2).

**Figure 2.**
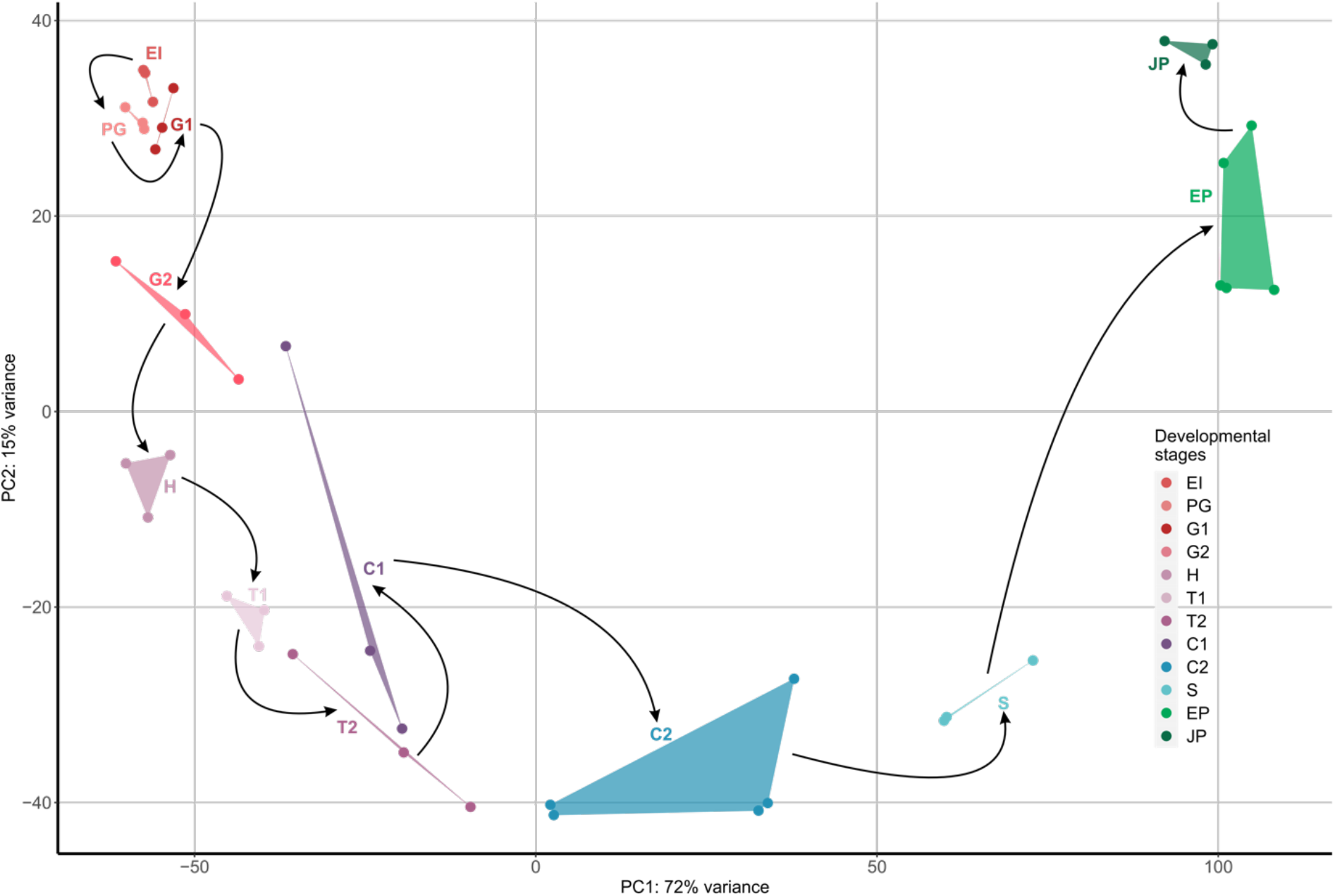
Principal component analysis (PCA) of transcriptomes shows time-resolved organization of *Vitis vinifera* somatic embryogenesis. *V. vinifera* developmental stages are shown in different colors, where shades of red represent early (EI – G2), shades of purple represent mid (H – T1), T2 and C1, and shades of green represent the late developmental stages (C2 – JP). Replicates are in the same color and connected with lines. Black arrows correspond to the experimental timeline of *V. vinifera* development that starts with EI and ends at JP. Developmental stages: early induction (EI), preglobular stage (PG), globular stage 1 (G1), globular stage 2 (G2), heart stage (H), torpedo stage 1 (T1), torpedo stage 2 (T2), cotyledonary stage 1 (C1), cotyledonary stage 2 (C2), seedling (S), seedling with epicotyl (EP) and juvenile plant (JP).

### SE ontogeny-phylogeny correlations

To determine whether *V. vinifera* somatic embryogenesis shows any correlation with the evolutionary trajectory of the plant lineage, we linked the transcriptome expression values of 12 SE developmental stages with the evolutionary age of *V. vinifera* genes and calculated the transcriptome age index (TAI) (Domazet-Lošo and Tautz 2010b) (Figure 1c; Supplementary Data 5). We assessed the evolutionary age of *V. vinifera* genes by phylostratigraphic approach (Domazet-Lošo et al. 2007, Futo et al. 2021) using a consensus phylogeny, which starts at the origin of cellular organisms and ends in *V. vinifera* as a focal species, and a large collection of reference genomes (see Methods, Supplementary Figure 3, Supplementary Data 6-7). We found that the TAI profile of *V. vinifera* development via SE has a pronounced, and statistically strongly supported, hourglass shape (Figure 1c). The evolutionary younger transcriptomes are predominantly expressed during early development (early induction; EI and preglobular stage; PG), after which increasingly older transcriptomes are recovered with the oldest estimates at the heart stage (H) (Figure 1c). As the mid-development advances, evolutionary younger transcriptomes start to be expressed again, with the peak at cotyledonary stage 2 (C2), which showed overall the evolutionary youngest transcriptome (Figure 1c). Finally, post-embryonic development stages, which come after cotyledonary stage 2 (C2), showed again a reverse trend with increasingly older transcriptomes (Figure 1c).

To test the stability of the recovered hourglass TAI profile we repeated the phylostratigraphic analysis using a range of e-value cutoffs (10 to 10^-40^) and recalculated TAI profiles (Futo et al 2021, Čorak et al. 2023). This robustness test which intentionally inflates false-positive and false-negative rates showed the stability of the hourglass TAI profile in the full range of tested e-value cutoffs (Supplementary Figure 4). This demonstrates that the TAI hourglass pattern of *V. vinifera* SE development is underpinned by a strong macroevolutionary imprint, which is resilient to the changes in e-value thresholds. The strength of these macroevolutionary signals prompted us to look more closely at how different phylogenetic levels (phylostrata) contribute to the overall TAI profile. By sequentially including genes from successive phylostrata, starting from the ps1 (Cellular organisms), we recalculated a set of TAI profiles and found that the clearly recognizable, and statistically significant, hourglass pattern is detectable from the origin of Diaphoretickes (ps8) (Supplementary Figure 5). These results suggest that the hourglass-shaped ontogeny-phylogeny correlations represent an ancient macroevolutionary imprint deeply embedded in the lineage that led to the origin of land plants.

To better understand the expression of genes from different phylostrata during *V. vinifera* SE, we conducted a relative expression analysis (Domazet-Lošo and Tautz 2010b, Futo el al. 2021). The genes that could be traced to the origin of cellular organisms (ps1) showed expression peaks at the heart stage (H) and in the juvenile plants (JP) (Supplementary Figure 6). Similarly, genes that originated during archaeal diversification (ps2-ps5) and eukaryogenesis (ps6) also peaked around the heart stage (Supplementary Figure 6). These expression peaks of evolutionary ancient genes at the heart stage (H) and in the juvenile plants (JP), explain in part why evolutionary oldest transcriptomes, as estimated by TAI analysis (Figure 1c), are expressed at these stages.

With the exception of Diaphoretickes-specific genes (ps8) that show the highest expression at the cotyledonary 2 stage (C2), genes that emerged in the period from the origin of Excavata/Diaphoretickes (ps7) till the origin of Streptophyta (ps11) also showed maximal expression in the heart stage (H) (Supplementary Figure 7a and b). This pattern demonstrates that genes that accompanied the early steps of the plant lineage diversification (ps7-ps11), which was unfolding in the aquatic environment, play an important role in the heart stage of extant somatic embryogenesis. In contrast, the genes that originated from the origin of Embryophyta (ps12) to the origin of Magnoliophyta (ps14) showed very dynamic regulation across SE development (Supplementary Figure 7c). Although genes from these evolutionary periods also have relatively high expression levels at the hearth stage (H), we detected strong additional peaks at the globular (G1 and G2), cotyledonary (C1 and C2), torpedo (T1 and T2), and seedling (S) stages (Supplementary Figure 7c). Together this pattern showed that the genes that originated during the early evolution of land plants (ps12-ps14) play an important role in the period from the globular stages (G1) to the seedling (S) stage, i.e., the central part of SE ontogeny (Figure 1a, Supplementary Figure 7c).

The evolutionary young genes that originated in the period from the origin of Eudicots (ps15) to the origin of focal species *V. vinifera* (ps18) cumulatively follow the hourglass pattern (Supplementary Figure 7d). Their upregulation is evident at the beginning of SE, in the early induction (EI) and pre-globular (PG) stages, as well as at the final phase of embryo maturation and during germination; i.e., in the cotyledonary 2 (C2) and seedling (S) stages (Supplementary Figure 7). Taken together, the SE developmental hourglass is underpinned by the upregulation of evolutionary older genes (Cellular organisms, ps1 to Spermatophyta, ps11) at the heart stage (H), and by the upregulation of evolutionary younger genes (Eudicots, ps15 to *V. vinifera*, ps18) at the beginning and the end of somatic embryogenesis (Supplementary Figures 6 and 7).

The TAI analysis relies on the evolutionary origin of unique sequences in the protein sequence space; hence it reflects a deep macroevolutionary history. However, to answer the question of whether the hourglass profile is maintained in the more recent evolutionary periods, we estimated the divergence rates between orthologous coding sequences of *V. vinifera* and *V. arizonica* (Figure 1d, Supplementary Figure 2a and Supplementary Data 5) and linked these values with SE expression trajectories. This approach originally used the ratio between nonsynonymous and synonymous substitution rates (dN/dS ratio) to calculate the transcriptome divergence index (TDI), assuming that synonymous substitution rates are a proxy of neutral evolution (Quint et al. 2012). However, synonymous substitutions cannot be considered neutral when selection acts on synonymous sites, e.g., via the codon usage bias (Futo et al. 2021). To account for this effect, we previously devised transcriptome nonsynonymous divergence index (TdNI) and transcriptome synonymous divergence index (TdSI), which allowed us to independently study how divergence rates at nonsynonymous and synonymous sites correlate with expression levels (Futo et al. 2021).

We found that both the transcriptome nonsynonymous divergence index (TdNI) and the transcriptome synonymous divergence index (TdSI) in *V. vinifera* – *V. arizonica* comparison show a clear and statistically supported hourglass profile (Figure 1d, Supplementary Figure 2a and Supplementary Data 5). The genes with the lowest divergence rates, at both nonsynonymous and synonymous sites, are predominantly expressed at the heart stage (H). In contrast, the genes with the highest divergence rates are prevailingly expressed in the induction (IE) and pre-globular (PG) stages, at the onset of SE development, and in the cotyledonary 2 (C2) stage, at the end of embryogenesis (Figure 1d, Supplementary Figure 2a and Supplementary Data 5). Interestingly, TdNI and TdSI curves closely follow the TAI pattern, suggesting that similar forces operate at different evolutionary scales. Additionally, transcriptome codon bias index (TCBI) showed that in *V. vinifera* SE genes which are expressed during the heart stage (H) and in juvenile plants exhibit the strongest codon usage bias (Supplementary Figure 2b and Supplementary Data 5). Altogether, these results confirm the existence of an hourglass-shaped ontogeny-phylogeny correlation in SE development in relatively recent evolutionary history that spans *V. vinifera* – *V. arizonica* divergence.

The TAI profile that we detected in the SE of *V. vinifera* could be tentatively compared to the one previously found in the ZE of *A. thaliana* (Quint et al. 2012) (Figure 1e), in the part that covers embryogenesis *sensu stricto*, i.e., from the early induction (EI) to the cotyledonary 2 stage (C2). These profiles have rather similar shape (Figure 1e), with a notable difference that the SE of *V. vinifera* expresses evolutionary oldest genes at the heart stage, while the ZE of *A. thaliana* exhibits evolutionary oldest transcriptome at the subsequent torpedo stage (Quint et al. 2012). Although TAI patterns for some parts of ZE postembryonic development of *A. thaliana* are available (Drost et al. 2016), it is unreliable to compare them to the SE postembryonic development of *V. vinifera* because the sampled stages in these studies do not match (Figure 1e). For example, some ZE stages such as “mature dry seeds”, “imbibed seeds”, “seeds at *testa rupture*” and “radicle protrusion” (Drost et al. 2016), simply do not exist as a part of the SE seedless development. Nevertheless, similar to our study, this previous work also reports the existence of phylogeny-ontogeny correlations in postembryonic ZE development of *A. thaliana* (Drost et al. 2016). Interestingly, the postembryonic drop in TAI values that we detected in the SE of *V. vinifera* (Figure 1c), highly resembles the pattern of postembryonic development in animals which also shows a progressive drop in TAI values (Domazet-Lošo and Tautz 2010b).

### Functional trends

To test the functional grouping of upregulated genes in specific developmental stages, we performed the enrichment analysis of GO functional categories (plant subset) across SE development. We found that every stage of SE has a specific battery of enriched GO functions (Figure 3, Supplementary Data 8), which indicates that functional transitions along SE rely on extensive transcriptional regulation. Genes with unknown functions are enriched in all SE stages (Figure 3, Supplementary Data 8), except in the heart stage (H). This pattern is congruent with the fact that the heart stage expresses evolutionary the oldest transcriptomes (Figure 1c), and that functionally older genes are more often functionally studied (Supplementary Figure 8). On the other hand, it is striking that many unannotated genes have regulated expression along SE (Figure 3) and that most of them emerged during the diversification of land plants (ps12-ps18, Embryophyta to *Vitis vinifera;* Supplementary Data 8 and Supplementary Figure 8). This shows that our understanding of how embryonic development of land plants has evolved is markedly incomplete.

It is rather reassuring that the GO term ‘somatic embryogenesis’ (GO:0010262) was strongly and significantly enriched at the early induction (EI) stage, which marks the onset of SE (Figure 3, Supplementary Data 8). However, to get a deeper understanding of this functional enrichment, we plotted individual expression trajectories of six genes that contribute to this signal (Figure 4a). Interestingly, five of them show a clear trend with the highest expression in the early induction (EI) and preglobular (PG) stages, followed by increasingly lower expression levels toward juvenile plant (JP) stage (Figure 4a, Supplementary Data 4 and 9). Some of these genes, such as AGL15, FUS3 and IAA30, have *A. thaliana* homologues which are known to be important in promoting somatic embryogenesis (Horstman et al. 2017).

Although GO annotation datasets are rather useful in screening general functional tendencies, they are nevertheless incomplete when it comes to the precise functional annotation of individual genes. We thus plotted the expression trajectories of additional genes which are known from the literature to play an important role in somatic embryogenesis but lack this type of annotation in our GO dataset (Figure 4b). Similar to GO-derived analysis, we found that all *V. vinifera* homologs of SE-important *A. thaliana* genes, such as BBM, L1L, PLT2 and SERK (Horstman et al. 2017), showed high expression at the onset of SE followed by increasingly lower expression levels toward the juvenile plant (JP) stage (Figure 4b, Supplementary Data 4 and 17). This rather regular expression profile of many important SE genes qualifies them as useful markers for tracking SE in future studies, e.g., in single-cell RNAseq experiments.

Epigenetic regulation of gene expression plays an important role during phase transitions in the life cycles of plants (Horstman et al. 2017, Markulin et al. 2021, Zhang et al. 2018). In our analysis, we detected two significant enrichments for the GO term ‘epigenetic regulation of gene expression’ (GO:0040029), which suggests that the early induction (EI) stage and the heart (H) stage are especially important transition phases for epigenome reprograming in the SE of *V. vinifera* (Figure 3). Because many genes (Supplementary Data 8) contribute to these enrichments, we illustrated general trends by depicting expression profiles for four representative epigenetic regulators (Figure 4c). For example, methylase DRM2, which is responsible for *de novo* methylation (Smertenko and Bozhkov 2014), showed increased expression in the early induction (EI) stage as well as the heart (H) stage (Figure 4c, Supplementary Data 4 and 17). We found a similar pattern for VAL1 (Figure 4c, Supplementary Data 4 and 9) which is a transcriptional repressor involved in histone methylation (Horstman et al. 2017, Yuan et al. 2021). On the other hand, DNA methylase MET1, required for the maintenance of DNA methylation during replication, and DNA demethylase DME (Smertenko and Bozhkov 2014) showed the highest expression in the heart (H) stage (Figure 4c, Supplementary Data 4 and 9).

It was suggested that the regulation of stress response plays an important role during SE because various stress-related genes have elevated expression in somatic embryos (Jin et al. 2014, Ci et al. 2022, Wickramasuriya and Dunwell 2015, Mendez-Hernandez et al. 2019). Our GO function enrichment analysis revealed that the GO term ‘response to stress’ (GO:0006950) is indeed significantly enriched in many stages over SE ontogeny including G1, G2, H, T1, C1, C2, EP and JP stage (Figure 3, Supplementary Data 8). Interestingly, the GO term ‘abscission’ (GO:0009908), which also could be linked to stress responses, is strongly enriched at the early induction (EI) stage (Figure 3, Supplementary Data 8). The full list of genes which contribute to the enrichment of these terms and their profiles are available in Supplementary Data 8 and 9. As an example, we depicted WRKY40, which is a transcriptional repressor that functions in plant responses to pathogens and abiotic stresses within complex regulatory networks that include other WRKY genes (Chen et al. 2010). WRKY40 showed high expression in the middle period of *V. vinifera* somatic embryogenesis, from the globular stage 1 (G1) to the torpedo stage 1 (T1) (Figure 4d, Supplementary Data 4 and 9). In contrast, peroxidase PRX73, another stress-related gene, was highly expressed during post-embryonic development including seedling (S), seedling with epicotyl (EP), and juvenile plant (JP) stages (Figure 4d, Supplementary Data 4 and 9), where it likely has a role in controlling root hair growth by modulating cell wall properties (Marzol et al. 2022).

Of all considered developmental stages, the heart (H) stage showed the most unique functional profile with several enriched GO functional categories related to embryo development (Figure 3, Supplementary Data 8). This suggests that at the functional level, the heart stage is a critical period for embryonic development where the expressions of key embryogenic genes converge. Again, these functional enrichments were underpinned by many genes (Supplementary Data 8 and 9). To illustrate major trends, we thus sorted out two examples (Figure 4e). ABI3, one of the central regulators that initiate maturation in the heart stage of *Arabidopsis* ZE (O’Neill et al. 2019), showed a peak of expression in the heart stage (Figure 4e, Supplementary Data 4 and 17). Similarly, a multifunctional enzyme ACC1, which is known for its role in cotyledon morphogenesis in the heart (H) stage of zygotic embryos (Baud et al. 2003), also showed maximal expression at the heart (H) stage of SE (Figure 4e, Supplementary Data 4 and 9).

The late developmental stages (C2 to JP) showed functional enrichments related to photosynthesis such as ‘plastid’ (GO:0009536), ‘chloroplast’ (GO:0009507), ‘thylakoid’ (GO:0009579), ‘photosynthesis’ (GO:0015979) and ‘response to light stimulus’ (GO:0009416) (Figure 3; Supplementary Data 8). This period corresponds to the switch from growth in the constant dark to a ‘long day’ regime (Figure 1a), hence one might expect the activation of photosynthesis-related genes. To show common expression trends of these genes, we depicted LHCA and PGR5 genes as examples (Figure 4f). LHCA genes encode for thylakoid light-harvesting chlorophyll-binding proteins that have a vital role in photosystem I (Jansson 1999), while PGR5 is essential for photoprotection and cyclin electron transport around photosystem I, especially in acclimation to fluctuating environments (Munekage et al. 2002; Wu et al. 2021). All of these photosynthesis-related genes showed a common trend where their expression values continuously increase during post-embryonic development (Figure 4f, Supplementary Data 4 and 9).

**Figure 3.**
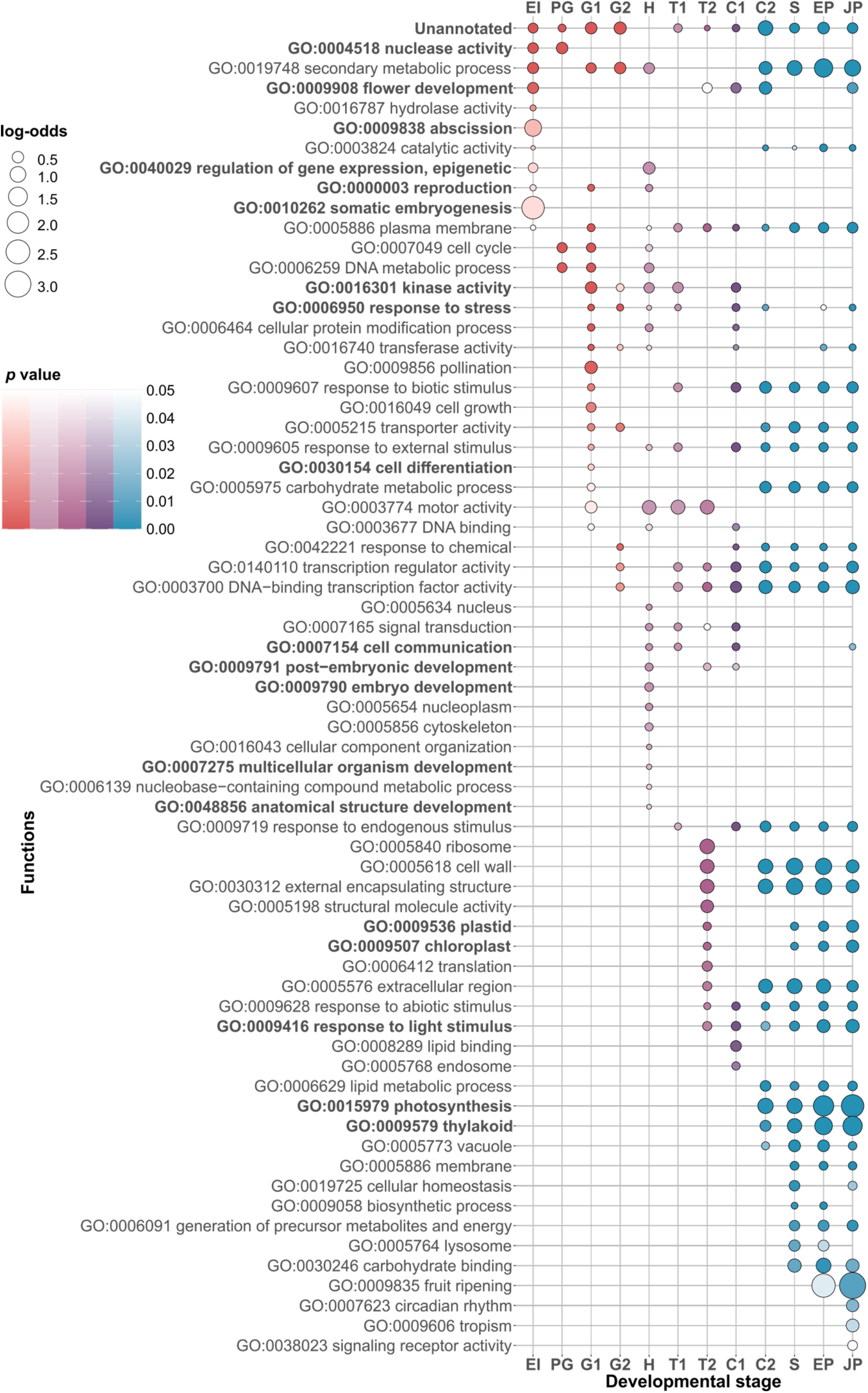
Functional enrichments along the somatic embryogenesis of *V. vinifera*. We analyzed the enrichments of GO functional categories in genes that are upregulated in the different stages of somatic embryogenesis. A gene was considered upregulated in a particular stage if it was transcribed 0.5 times (log_2_ scale) above the median of its overall transcription profile. The frequency of a GO annotation per stage is compared to the frequency of that annotation in the whole *V. vinifera* genome and shown as log-odds (bubble graph). The log-odds higher than zero denote that the frequency of annotation in a given developmental stage is higher than the expected frequency estimated from the whole genome. The significance of these functional enrichments was tested by two-tailed hypergeometric test. The *p*-values were adjusted for multiple testing (see Material and Methods). Only significant enrichments are shown. The color code follows expression phases in Figure 1a: early development (red), mid development (violet), late development (turquoise).

**Figure 4.**
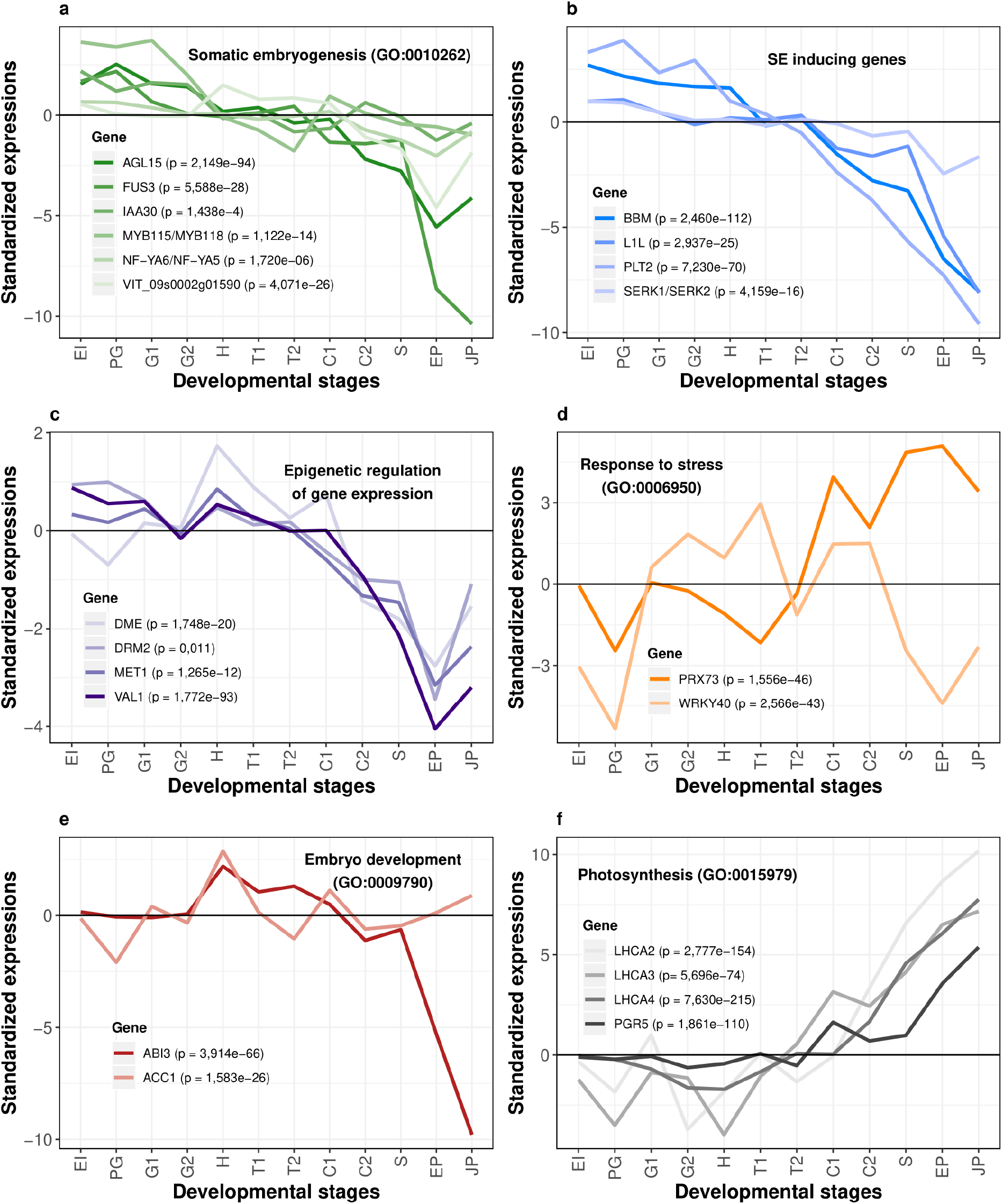
Standardized expression profiles of selected *V. vinifera* genes along somatic embryogenesis. **a)** Standardized expression profiles of genes annotated with GO term ‘Somatic embryogenesis’ (GO:0010262) that showed an enrichment signal in Figure 3 (EI stage). **b)** Selected genes that have an important role in the induction of somatic embryogenesis according to the literature. **c)** Three representative genes (DME, DRM2, MET1) annotated with GO term ‘Epigenetic regulation of gene expression’ (GO:0040029). This GO term showed an enrichment signal in Figure 3 (EI and H stages). VAL1 is described in the literature to be important for chromatin modification. **d)** Two representative genes (PRX73, WRKY40) annotated with GO term ‘Response to stress with a signal’ (GO:0006950). This GO term showed enrichment signals in Figure 3 (G1, G2, H, T1 and C1 stages). **e)** Two representative genes (ABI3, ACC1) annotated with GO terms ‘Embryo development’ (GO:0009790), ‘Multicellular organism development’ (GO:0007275) and ‘Anatomical structure development’ (GO:0048856). These GO terms showed enrichment signals in Figure 3 (H stage); **f)** Four representative genes (LHCA2, LHCA3, LHCA4, PGR5) annotated with GO terms ‘Plastid’ (GO:0009536), ‘Chloroplast’ (GO:0009507), ‘Response to light stimulus’ (GO:0009416), ‘Photosynthesis’ (GO:0015979) and ‘Thylakoid’ (GO:0009579). These GO terms showed enrichment signals in late developmental stages (C2, S, EP, JP) in Figure 3. Differential expression along SE were tested by LRT test as implemented in *DESeq2* (Love et al. 2014). Resulting *p-*values corrected by FDR are shown for every gene. Standardized expression value of 0 (black horizontal line) represents the median of expression levels for a respective gene. Gene names were obtained by searching for *V. vinifera – A. thaliana* orthologs in TAIR database (Berardini et al. 2015). Correspondence between *V. vinifera* and *A. thaliana* genes together with standardized gene expression values can be found in Supplementary Data 4, and standardized gene expression profiles in Supplementary Data 9.

## Discussion

Somatic embryogenesis as an experimental model has the advantage over zygotic embryogenesis because it enables production of genetically identical somatic embryos in large numbers. On the other hand, unsynchronized development of somatic embryos, as well as their aggregation into physically compact clusters represent the main obstacles of current SE protocols. Both problems limit the isolation of individual developmental stages without embryo wounding. For example, despite its huge advantages as a model system, *Arabidopsis* somatic embryos are fused along their contact surfaces. This leads to the formation of unsynchronized embryo clusters that cannot be easily separated without tissue damage (Gaj 2001, Kadokura et al. 2018). Another limitation of *Arabidopsis* SE is the low proportion of embryos that complete embryogenic development, which consequently leads to the low frequency of plantlet regeneration. This limitation is further provoked by the culturing of somatic embryos for long time (Bhatia and Bera 2015). To address these issues, here we developed a comparatively rapid protocol with a rather low input of growth regulators that enables synchronized development of unfused individual embryos with high plantlet regeneration potential. We see our *V. vinifera* ‘Malvasia Istriana’ SE induction system as a highly reproducible and potentially widely applicable model for plant SE research.

Plant embryogenesis is an old process that most likely emerged at the root of Embryophyta (ps12), predating the later invention of seeds and flowers in seed plants (Schneider 2018, Radoeva et al. 2019, Rensing and Weijers 2021). In this context, zygotic embryogenesis in flowering plants could be considered an evolutionary derived process, which includes innovations such as endosperm formation, desiccation, and dormancy (Linkies et al. 2010, Radoeva et al. 2019). In contrast, somatic embryogenesis, which does not depend on these adaptations, might be a better representation of the ancestral embryogenic trajectory of land plants.

Moreover, somatic embryogenesis is not limited to seed plants, as this process also exists in ferns, which seem to show higher potential for SE induction than spermatophytes (Mikula et al. 2015). This indicates that probably all clades of Embryophyta retained the potential for somatic embryogenesis. Taken together, the hourglass pattern that we discovered in the somatic embryogenesis of *V. vinifera* is most likely a better proxy of ancestral phylogeny-ontogeny correspondence that underpinned Embryophyta diversification, than the one described in the zygotic embryogenesis of *Arabidopsis* (Quint et al. 2012).

The original study that discovered hourglass-shaped correlations between phylogeny and ontogeny in *Arabidopsis* ZE predicted that the phylotypic stage (the waist of hourglass) in plants should be placed somewhere between the globular and the heart stage (Quint et al. 2012). This prediction assumes that a phylotypic stage should possess all major body parts at their final anatomical position in the form of undifferentiated cell aggregates (Quint et al. 2012). However, this study detected a disparity between this prediction and the recovered phylotranscriptomic profile that shows the waist at the subsequent torpedo stage — a developmental stage which marks the beginning of the maturation phase linked to seed formation (Quint et al. 2012). Obviously, this discrepancy between the theoretical predictions and the actual pattern is not present in the somatic embryogenesis of *V. vinifera* where the waist of hourglass phylotranscriptomic profile is placed at the heart stage, as originally expected. This finding suggests that somatic embryogenesis, beside many technical advantages (Radoeva et al. 2019), is a better model to study general developmental principles in land plants than zygotic embryogenesis.

From this perspective, somatic embryogenesis could be viewed as an atavistic trait (Bonet et al. 1998). Although in some plants, like in some species of the genus *Kalanchoe*, SE is an integral part of the life cycle, in many others it can only be activated upon stress induction (Garcês et al. 2007). It seems that atavistic characters in plants are generally induced by the impact of stress (Bonet et al. 1998), which occasionally pushes plant cells to the expression of ancient developmental programs (Bonet et al. 1998, Jin et al. 2014, Mendez-Hernandez et al. 2019). In some instances, the cooption of atavistic programs obviously has an adaptive value. A good example is the case of constitutive plantlet-forming species in the genus *Kalanchoe*, where coopting somatic embryogenesis into leaves rescues the propagation potential of *Kalanchoe* species that possess nonviable seeds (Garcês et al. 2007). However, whether the stress induction of somatic embryogenesis in natural settings has a broader prevalence and adaptive value remains unclear.

Plants and animals are intrinsically multicellular organisms that independently evolved their multicellularity (Niklas and Newman 2020, Domazet-Lošo et al. 2024). Yet, their embryogenesis shows remarkable system-level analogy in the form of hourglass-shaped phylogeny-ontogeny correlations (Quint et al. 2012, Drost et al. 2017) and in the macroevolutionary dynamics of genome complexity change (Domazet-Lošo et al. 2024). We found here that analogies in phylogeny-ontogeny correleations are especially pronounced if animal development is compared to somatic embryogenesis in plants. This similarity primarily relates to the positioning of the phylotypic stage in the mid-embryogenesis where the primordia of all major body parts are placed at their final anatomical positions (Domazet-Lošo and Tautz 2010b, Quint et al. 2012). However, it is also indicative that we found comparable trends in later phases of ontogeny. Namely, with the formation of seedling (S) we observed that *V. vinifera* plantlets increasingly express older and less diverged transcriptomes. This strongly resembles the pattern in animals where aging adult animals express increasingly older genes (Domazet-Lošo and Tautz 2010b).

However, it remains unclear which evolutionary forces govern these analogies. In animals, several studies tried to resolve evolutionary mechanisms which maintain the developmental hourglass (Zalts and Yanai 2017, Hu et. al. 2017, Liu et al. 2021, Uesaka et al. 2022). The full picture has not been revealed yet, but it seems that a combination of purifying selection (Zalts and Yanai 2017) linked to pleiotropic effects at mid-embryogenesis (Hu et. al. 2017) and positive selection acting on the early and late phases of embryogenesis (Liu et al. 2021) shape the hourglass profile in animals. Similar studies in plants are currently lacking; however, there is a possibility that evolutionary mechanisms behind developmental hourglass in plants are more complex than in animals. Namely, a recent study found that mutations occur less frequently in functionally constrained *A. thaliana* genome regions (Monroe et al. 2022). If this finding stands the test of time (Wang et al. 2023, Monroe et al. 2023), this would open the possibility that developmental hourglass in plants, in addition to purifying and positive selection, is underpinned by mutational bias. In any case, the difference in the relative positioning of the hourglass waist between zygotic and somatic embryogenesis that we found in this work implies that the phylotypic stage can be subject to heterochronic changes.

In sum, we conclude that macroevolutionary imprint, in the form of hourglass-shaped ontogeny-phylogeny correlations, is deeply hardwired in plant ontogeny and is largely resilient to alternative developmental routes, such as zygotic and somatic embryogenesis. Our discovery that the shape of ontogeny-phylogeny correlations in somatic embryogenesis better fits with theoretical expectations, and that it more closely resembles analogous patterns in animals, suggests that somatic embryogenesis is likely a primordial embryogenic program in plants.

## Methods

### Plant Material

Inflorescences of *Vitis vinifera* L. ‘Malvasia Istriana’ (Malvazija istarska) were acquired from the National Collection of Autochthonous Grape Varieties of the University of Zagreb, Faculty of Agriculture experimental station “Jazbina” during the May/June of 2017, which was approximately 2-3 weeks before anthesis. Alternatively, we induced flowering by placing the basal part of dormant vine cuttings (approximately 30 cm in length) into distilled water and by exposing them to 24 °C and 16/8 photoperiod using daylight florescent tube (40 W, 400-700 nm, 17 W/m^2^). Anthers were isolated from the buds of sterilized inflorescences according to the procedure described in Malenica et al. (2020).

### Induction of embryogenesis

Modified MS medium (Murashige and Skoog 1962), lacking glycine and with MS-nitrogen sources substituted with X6 nitrogen sources (Li et al. 2008), were used as a basic medium in this study. Induction medium was based on the basic medium which in addition contained 5 µM BAP (6-benzyladenine), 2.5 µM 2,4-D (2,4-dichlorophenoxyacetic acid), 2.5 µM NOA (naphthoxyacetic acid) (Dhekney et al. 2009), sucrose (2 % w/v) and agar (7 % w/v). The pH of media was adjusted to 5.8 before sterilization at 121 °C, 103 kPa for 15 minutes.

Whole flower buds were aseptically removed from the inflorescence and opened by cutting the basal side of the bud. Filaments were excised at their bases using a medical needle under the stereomicroscope and together with attached anthers placed on the medium with their adaxial side facing the surface. Between 20-25 explants were cultivated in 30 x 10 mm Petri dish at 24°C in the dark.

### Embryo maturation

Globular embryos were transferred separately onto the hormone-free basic medium suitable for somatic embryo development, with addition of 0.5 g/L activated charcoal (Li et al. 2008). Cultures were cultivated at 24 °C in the dark.

### Somatic embryo germination and plant regeneration

Cotyledonary stage embryos developed on hormone-free basic medium were induced to germinate on the embryo germination medium (EG) supplemented with 10 µM IAA and 1 µM GA3 (López-Pérez et al. 2005). Cultures were exposed to 24 °C and 16/8 photoperiod using daylight florescent tube (40 W, 400-700 nm, 17 W/m^2^). The details of this procedure are described in Malenica et al. (2020).

### Selection of different developmental stages of somatic embryos

Classification and selection of each specific developmental stage during and post-embryogenesis were based on morphological criteria described previously for seven *Vitis vinifera* cultivars (Malenica et al. 2020; Figure 1a). The yellowish proembryogenic masses (EI) were formed on the filament tip. To collect single cells and cell clusters, the proembryogenic masses were mechanically separated from the filament and cultivated in liquid induction medium for 16 h with constant agitation in the growth chamber at 24 °C in the dark. After sieving the cell suspension through a 150 µm nylon mesh, cells and small cell clusters from the liquid phase were collected on 50 µm nylon mesh and split in two portions. One half was recultured for testing the embryogenic competence (induction success), while the rest was shock-frozen in liquid nitrogen and stored at -80 °C until further use. Only if recultured tissues were efficient in the embryo production (preglobular and globular embryo formation within 2 weeks), the corresponding frozen sample was used further.

Preglobular (PG) and globular stage (G1 and G2, different in size) embryos were isolated from the embryogenic tissue by sieving the tissue through the metal mesh to remove the older stages and large clusters. The filtrate fraction that contained mostly PG and G embryos was washed further with a fresh liquid basic medium by using a 150 µm nylon mesh to remove single cells and small clusters.

The PG embryos were distinguished from the G stage according to the morphology of the epidermal cell layer. In contrast to well-formed epidermis of discrete globular stage embryos, the preglobular stage was mainly attached to the explant tissue. In the cases when they were detached from the explant tissue we detected them by their surface which was not smooth and even (Figure 1a). After collecting each stage separately, they were again re-washed with basic medium using a 150 µm nylon mesh.

Later embryogenic stages (heart H, torpedo T1 and T2, cotyledonary C1 and C2) were isolated using fine forceps and a needle, based on their specific shapes and sizes observed under a binocular microscope. Collected embryos were washed with basic medium by using a 150 µm nylon mesh to remove the remaining tissue. The stages of postembryogenic development were determined according to the development of root hairs, epicotyl, and the first pair of leaves.

### RNA Extraction

Total RNA was isolated from somatic embryos using the RNeasy Plant Mini kit (Qiagen, Hilden, Germany) with slight modification of the manufacturer’s protocol. Depending on the developmental stage, samples contained between 50 and 70 individual embryos. Each sample was homogenized in 450 μl RLT buffer with 20 mg PVP (polyvinylpyrrolidone; Sigma, St. Louis, USA) in a 2 mL plastic tube using four stainless steel beads (3 mm diameter). The bead beater (Retsch MM200, Haan, Germany) was set to 30 Hz for 3 min. The homogenate was filtered in a Qiashreader column at 10,000 rpm for 1 min. The supernatant was mixed with 0.5 volumes of 96% EtOH, transferred to an RNA binding column, and centrifuged according to the manufacturer’s instructions. DNA removal was performed by applying 80 μl of DNase I working solution (10 μL DNase stock + 70 μL 1x RDD buffer; RNase-Free DNase Set, Qiagen, Hilden, Germany) to the column and incubated for 15 min at room temperature. Then 350 µL of RW1 buffer was added to the column and centrifuged at 10,000 rpm for 15 s. This washing step was repeated one more time. Further, two washing steps were performed with 500 μL of RPE buffer at 10,000 rpm for 15 s. The RNA was eluted with 40 μL of Tris-HCl pH 6.8 (Ambion, Austin, USA). Finally, 1 μl of RNase inhibitor (40 U/μL; Thermo Scientific, Waltham, USA) was added to the sample and incubated for 5 min at room temperature and stored at -20 °C before sequencing.

Isolated RNA was quantified on a Nanodrop spectrophotometer (ThermoFischer, Waltham, USA). The A260/280 values for all samples were between 1.8 and 2.0 and the RNA concentrations were in the range of 20-190 ng/µL, depending of the embryo developmental stage. The RNA quality was tested by 1 % agarose gel electrophoresis (1xTAE).

### RNA Sequencing

Total RNA extracted from each somatic embryo developmental stage (EI to JP) was sent to EMBL Genomics Core Facility (Heidelberg, Germany) for quality check, rRNA depletion, cDNA library preparation and high throughput sequencing. The samples were sequenced in five (C2 and EP stages) and tree replicates (the remaining 10 stages) using Illumina NextSeq 500 platform (read length 75 bp, paired-end). Sequence quality and read coverage were checked using the FastQC V0.11.9 (Andrews 2010) with a satisfactory outcome for each of the samples. In total, 3,945,355,234 paired-end sequences (75 bp) were mapped onto the *V. vinifera* reference genome (NCBI Assembly Accession: 12X, GCA_000003745.2) using BBMap V38.75 (Bushnel 2014) with an average of 95.48% of mapped reads per sample (Supplementary Data 1). On average, we mapped 90 million reads per replicate (Supplementary Data 1). Mapping was performed using the standard settings with the option of trimming the read names after the first white space enabled. Generating, sorting, and indexing of BAM files was done by using SAMtools V1.11 (Li et al. 2009). These files were then used for the downstream data analyses in R V4.0.4 (R Development Core Team 2008) using custom-made scripts. Briefly, quantification of mapped reads for each *V. vinifera* open reading frame was done using the R *rsamtools* package V2.10.0 (Morgan et al. 2021) and raw counts for 29,839 (out of 29,971) open reading frames were retrieved using the *GenomicAlignments* R package V1.30.0 (Lawrence et al. 2013). We estimated expression similarity between replicates and developmental stages using the PCA analysis (Figure 2) implemented in the R package *DESeq2* V1.34.0 (Love et al. 2014). The obtained results were visualized in the R package *ggplot2* V3.3.5 (Wickham 2016).

### Transcriptome Analysis

To prepare the raw count values for the subsequent analysis, we normalized them by calculating the fraction of transcripts (τ) (Li et al. 2010). The reasoning behind using τ for downstream calculation of evolutionary measures has been discussed in previous work (Domazet-Lošo and Tautz 2010b, Conesa et al. 2016, Futo et al. 2021). We resolved the replicates by calculating the replicate median for each developmental stage. The obtained normalized transcript expression values were used to calculate evolutionary indices (Supplementary Data 5), and relative expression values of phylostrata.

Following a pipeline introduced in previous work (Futo et al. 2021), we calculated the standardized expression values of each gene for use in GO enrichment analysis (Figure 3), clustering (Supplementary Data 3 and 4), and profile visualization (Figure 4, Supplementary Data 4 and 9). Briefly, we discarded genes that had the expression value of zero in more than two developmental stages, removing 3,790 genes from the dataset. For genes that had a single stage with the expression value of zero, we interpolated it with the mean of the two adjacent stages (1,064 genes), or if the expression value of zero was in the first or last stage, we transferred the value of the only neighboring stage directly (390 genes). Lastly, the expression values for each gene were normalized to the median and log_2_ transformed, resulting in the standardized expression values for 26,181 genes.

The standardized expression profiles were visualized (Figure 4, Supplementary Data 3 and 4) using the R package *ggplot2* V3.3.3 (Wickham et al. 2016). Genes selected for expression profile visualization in Figure 4 were selected based their GO annotations (Figure 4a, c, d, e and f) and orthology to SE-relevant *A. thaliana* genes (Horstman et al. 2017) (Figure 4b and c). Gene names and gene orthologs between *A. thaliana* and *V. vinifera* were selected based on TAIR database (Berardini et al. 2015).

To cluster a large dataset with 26,181 genes, we first split the dataset into 13 randomly sampled groups of genes; 12 groups consisted of 2,015 genes and one of 2,014 genes. Using the DP_GP_cluster (McDowell et al. 2018) with the maximum Gibbs sampling iterations set to 500, we clustered the standardized expression profiles of genes within each of the 13 groups which yielded 1,157 gene clusters in total. For each of these clusters, we calculated the mean standardized expression profile. Using again the DP_GP_cluster with the maximum Gibbs sampling iterations set to 500 we clustered these 1,157 mean standardized expression profiles, which finally resulted in the 85 clusters composed of 26,181 genes (Supplementary Data 3 and 4).

We tested the transcriptome similarity between different developmental stages by calculating Pearson’s correlation coefficients (*R*) using standardized expression values for all-against-all comparisons and visualizing it on a heatmap (Figure 1b). Using a pipeline implemented in the *DESeq2* V1.30.1 (Love et al. 2014) R package, we estimated the pairwise differential gene expression between the individual developmental stages (Supplementary Data 2 and Supplementary Figure 1), as well as overall differential expression for every gene across all developmental stages (Supplementary Data 2) with the likelihood ratio test (LRT) implemented in the same package.

### Functional Enrichment Analysis

Due to the lack of a comprehensive set of functional gene annotations, we used eggNOG-mapper V2.0 (Cantalapiedra et al. 2021) to annotate the *V. vinifera* genome. We obtained the best annotation data using the default search filters and limiting the taxonomic scope to Eukaryota. This resulted in 18,425 genes annotated with GO annotations (Ashburner et al. 2000) (Supplementary Data 8). We then performed the functional enrichment of individual developmental stages using the assigned GO annotations. To simplify analyses, we limited GO terms used in the functional enrichment to the GO Plant subset downloaded from the Gene Ontology Resource website (GO version: 10.5281/zenodo.4735677, May 20, 2021). In addition, we included in this GO Plant subset the missing term GO:0010262 “Somatic embryogenesis” because it was relevant for our research. We tested enrichment of these GO terms in each developmental stage for a set of genes that had in that particular stage a standardized expression value of at least 0.5 (log_2_ scale) above the median of their overall expression profile across SE (Figure 3, Supplementary Data 4). All enrichment analyses were performed using the two-way hypergeometric test (Supplementary Data 8). To adjust for multiple comparisons, we corrected the *p* values using the Benjamini and Hochberg procedure (Benjamini and Hochberg 1995).

### Evolutionary Measures

The phylostratigraphic procedure was performed as described in previous work (Domazet-Lošo et. al. 2007; Domazet-Lošo and Tautz 2010a). Following the latest phylogenetic literature (Leliaert et al. 2012; Morris et al. 2018), we constructed a consensus phylogeny covering the divergence from the last common ancestor of all cellular organisms to *V. vinifera* as the focal organism (Supplementary Figure 3, Supplementary Data 6-7). Phylogenetic trees were visualized and annotated in the iTOL v6 online tool (Letunić and Bork 2021) (Supplementary Figure 3 and Supplementary Data 6). The choice of internodes (phylostrata) in the consensus phylogeny depended on their phylogenetic support in the literature, the availability of reference genomes for the terminal taxa, and their importance for evolutionary transitions.

We retrieved the full set of protein sequences for 427 terminal taxa, five from NCBI and 422 from Ensembl database (Supplementary Data 7). We prepared the referent protein sequence database for sequence similarity searches by checking the files for any inconsistencies, adding taxon tags to the sequence headers of all sequences, and leaving only the longest splicing variant of each eukaryotic gene. The phylostratigraphic map of *V. vinifera* was constructed by comparing 29,927 *V. vinifera* protein sequences against the referent protein sequence database using blastp algorithm V2.9.0 (Altschul et al. 1990) with the e-value threshold of 10^-3^. Discarding all protein sequences which did not return a significant match left us with 29,623 protein sequences in the sample. We then mapped those 29,623 protein sequences on the 18 internodes (phylostrata) of the consensus phylogeny (Supplementary Data 7). Each protein sequence was assigned to the oldest phylostratum where it still had a blast hit (Domazet-Lošo et. al. 2007; Domazet-Lošo and Tautz 2010a).

For each developmental stage, using the expression values of 29,623 protein-coding genes, we calculated the transcriptome age index (TAI) (Figure 1c, Supplementary Data 5):

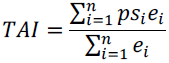

where *ps_i_* is an integer that represents the phylostratum of the protein *i*, *e_i_* is the normalized expression value of the gene *i* and *n* is the total number of genes analyzed. Previous work discussed the biological interpretation of TAI and its statistical properties at length (Domazet-Lošo and Tautz 2010b).

To compare *V. vinifera* SE TAI profile to *A. thaliana* ZE TAI profiles (Figure 1e), we downloaded previously obtained *A. thaliana* ZE TAI values (Quint et al. 2012). The TAI values from SE and ZE were normalized as follows:

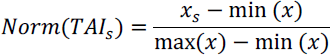

where *Norm*(TAI_s_) represents the normalized TAI value for the developmental stage *s*, *x*_s_ is the unnormalized TAI value of a developmental stage s, while min(*x*) and max(*x*) are the minimum and maximum unnormalized TAI values across all developmental stages. The obtained normalized TAI values were plotted on the Y axis in a range from 0 (lowest TAI value) and 1 (highest TAI value) (Figure 1e).

To test the robustness of the TAI profile and the phylostratigraphic pipeline in general, we used the blastp algorithm V2.9.0 (Altschul et al. 1990) to construct additional phylostratigraphic maps with different e-value cutoffs (10, 1, 10^-1^, 10^-2^, 10^-3^, 10^-5^, 10^-10^, 10^-15^, 10^-20^, 10^-30^, 10^-40^) (Supplementary Data 7 and Supplementary Figure 5). To calculate the divergence rates of *V. vinifera* proteins, we used the pipeline available in the R package *orthologr* V0.4.0 (Drost et al. 2015). Using blastp reciprocal best hits with 10^-5^ e-value threshold, we found 18,761 orthologs in *Vitis arizonica* (Grape genomics: Vitis arizonica cl. b40-14 V1.1, 10.5281/zenodo.3827985) (Supplementary Data 5). After globally aligning *V. vinifera – V. arizonica* ortholog pairs using the Needleman-Wunsch algorithm, we used pal2nal to construct codon alignments (Suyama et al. 2006). We calculated the nonsynonymous substitution rates (d*N*), the synonymous substitution rates (d*S*), and the sequence divergence rates (d*N*/d*S*) using Comeron’s method (Comeron 1995).

For each developmental stage, using the d*N* values of 18,751 genes, we calculated the transcriptome nonsynonymous divergence index (TdNI) (Figure 1e, Supplementary Data 5):

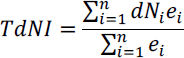

where d*N_i_* is a real number that represents the nonsynonymous divergence of gene *i*, *e_i_* is the normalized transcript expression value of the gene *i*, and *n* is the total number of genes analyzed (Futo et al. 2021). For each developmental stage, using the d*S* values of 18,727 genes, we calculated the transcriptome synonymous divergence index (TdSI) (Supplementary Data 5 and 14):

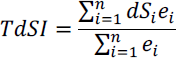

where d*S_i_* is a real number that represents the synonymous divergence of gene *i*, *e_i_* is the normalized transcript expression value of the gene *i*, and *n* is the total number of genes analyzed (Futo et al. 2021). TdNI and TdSI are weighted means of nonsynonymous and synonymous sequence divergence respectively. For each developmental stage, using the effective number of codons (ENC) measure (Wright 1990) for 18,727 genes, we calculated the transcriptome codon bias index (TCBI) (Supplementary Figure 2 and Supplementary Data 5):

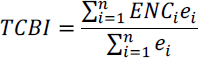

where ENC is a real number that represents the codon usage bias of gene *i*, *e_i_* is the normalized transcript expression value of the gene *i*, and *n* is the total number of genes analyzed (Futo et al. 2021). A lower TCBI value corresponds to a transcriptome with higher codon usage bias, and vice versa. To calculate the statistical significance of TAI, TdNI, TdSI, and TCBI profiles, we used flat-line test implemented in the R package *myTAI* V0.9.3 (Drost et al. 2018). The relative expression of genes for a certain phylostratum (ps) and developmental stage (s) (Supplementary Figure 6-7) were calculated as follows:

Where 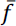 is the mean normalized expression value of genes from phylostratum (ps) in the given stage, while 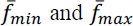 are the minimal and maximal mean normalized expression values of genes from the phylostratum (ps) across all stages (Domazet-Lošo and Tautz 2010b). Relative expression values for a certain phylostratum range from 1 in the developmental stage where the mean normalized expression value is the highest and 0 where the mean normalized expression value is the lowest.

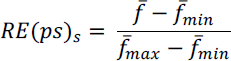

## Data availability

All transcriptome data have been deposited in NCBI’s Gene Expression Omnibus and are accessible through GEO Series accession number GSE234231.

## Acknowledgments

This work was supported by the City of Zagreb, the Croatian Science Foundation under the project IP-2016-06-5924, the Adris Foundation, and the European Regional Development Fund (KK.01.1.1.01.0009 DATACROSS) to T.D.-L.

## Author contribution

T.D.-L., D.L.-L. and N.M. initiated and conceptualized the study, D.L.-L., N.M. and M.J. collected the plant material and performed experiments, S.K., K.B.V., M.F., N.Č. A.T., N.K., M.D.-L., K.V., and T.D.-L. performed bioinformatic analyses, S.K., K.B.V., N.K., and T.D.-L. prepared the figures and tables for publication. S.K., D.L.-L., N.M. and T.D.-L. wrote the manuscript. All authors read and approved the manuscript.

---

**Supplementary Figure 1.**
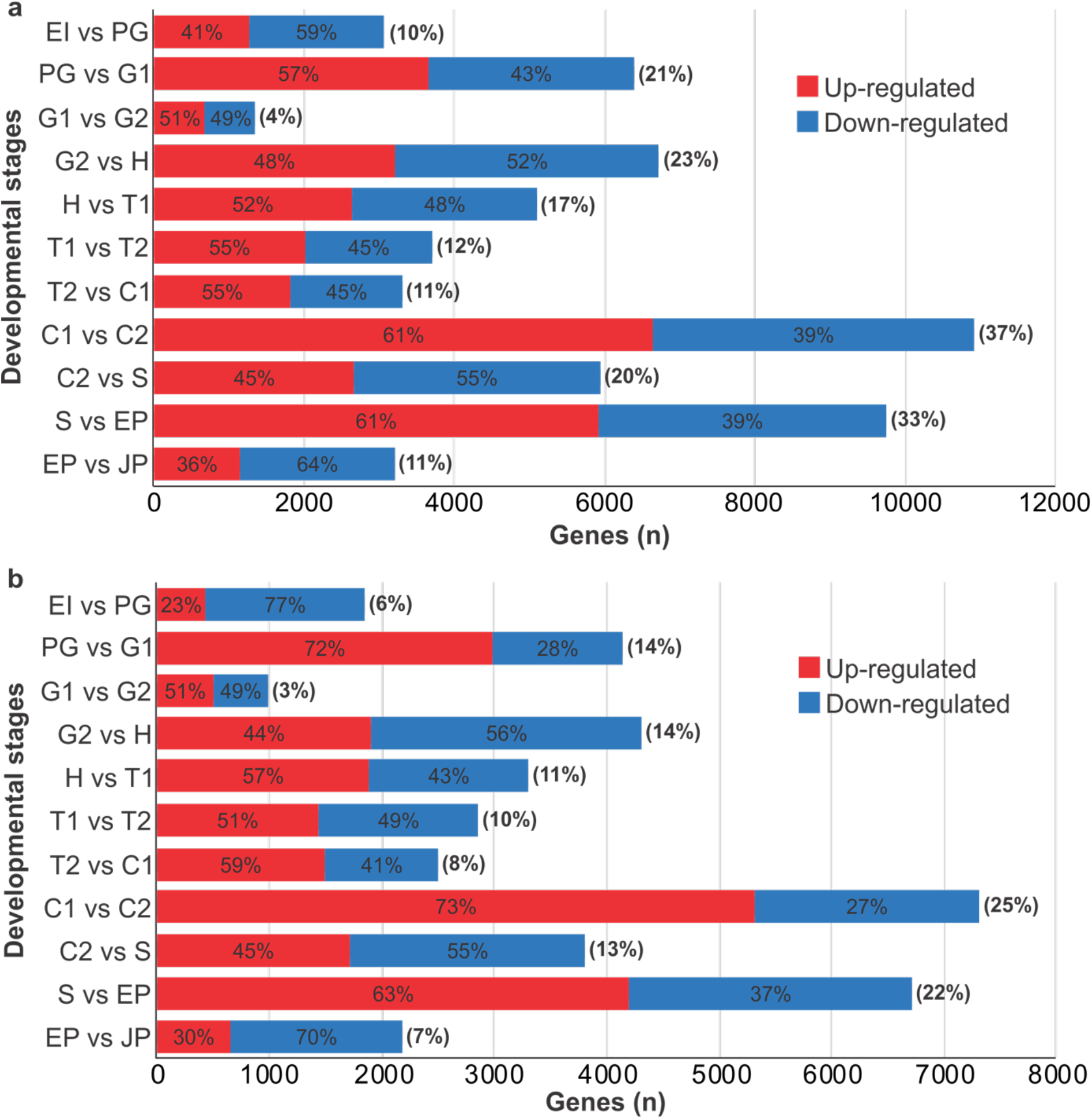
Differentially expressed genes between the successive developmental stages of *V. vinifera* somatic embryogenesis. The percentages in parenthesis show the percentage of differentially expressed genes between the two successive developmental stages calculated by comparison to all expressed genes (29,839). Red bars denote the percentage of up-regulated genes, while blue bars mark percentage of down-regulated genes between the two successive stages in reference to all differentially expressed genes during SE. We estimated pairwise differential expression using a pipeline implemented in the *DESeq2* R package (**a,** p-value < 0.05; **b,** p-value < 0.05 and fold change > 2).

**Supplementary Figure 2.**
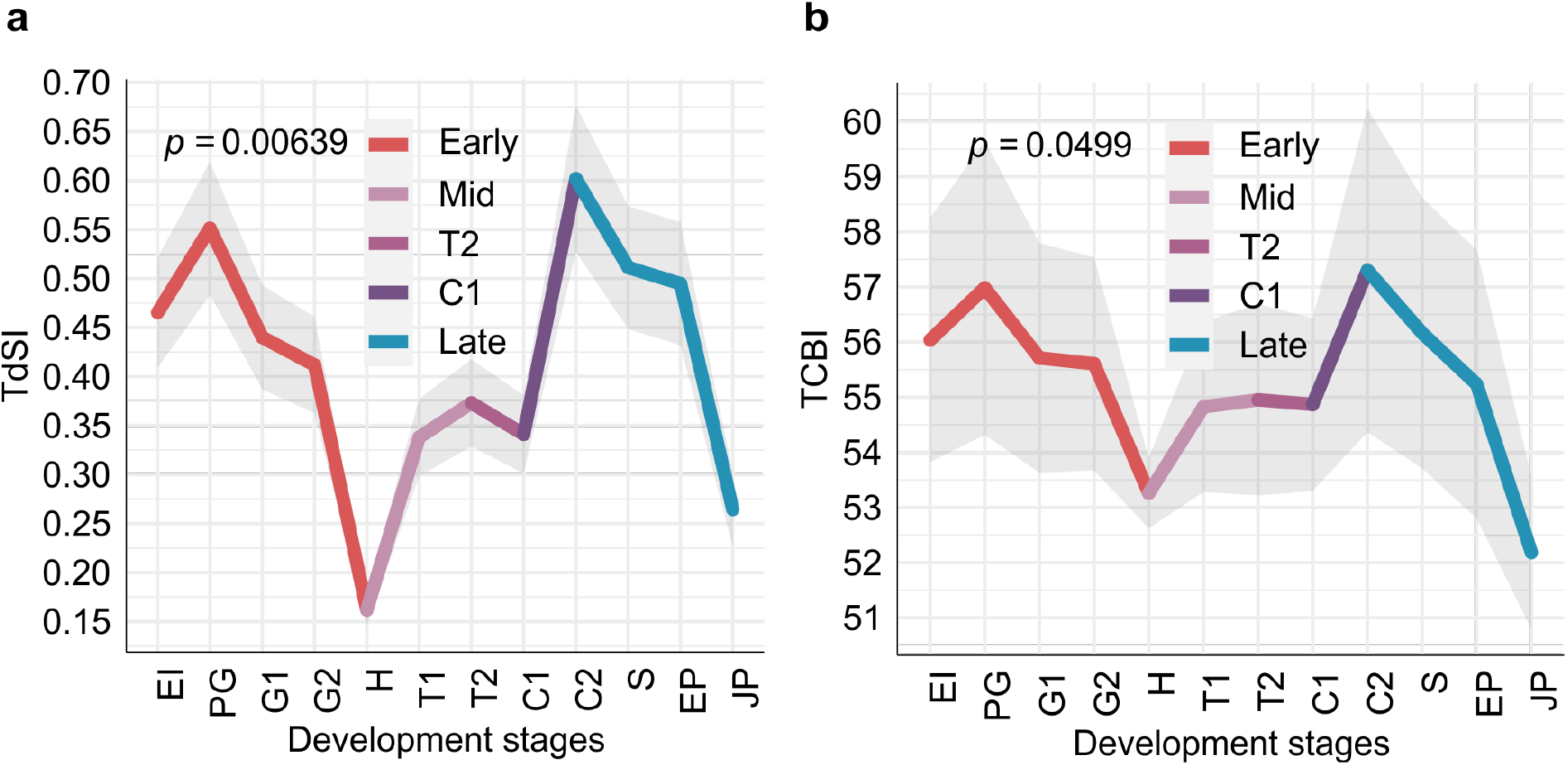
The synonymous divergence and codon bias exhibit an hourglass pattern. **a**, The transcriptome nonsynonymous divergence index (TdSI), and **b,** the transcriptome codon bias index (TCBI) profile exhibit statistically significant hourglass patterns, with genes conserved at synonymous sites and genes expressing strong codon usage bias preferentially expressed during mid-development. Divergence rates for TdSI were estimated by *V. vinifera – Vitis arizonica* comparison (see Material and Methods). Codon usage bias was estimated for *V. vinifera* genes using ENC measure (see Material and Methods). The *p* values were calculated using the flat line test while the grey shaded area represents ± one standard deviation estimated using permutation analysis (see Material and Methods). Periods of similar gene expression within the somatic embryogenesis are color-coded: early (red), mid, T2, and C1 (different shades of purple), and late (blue).

**Supplementary Figure 3.**
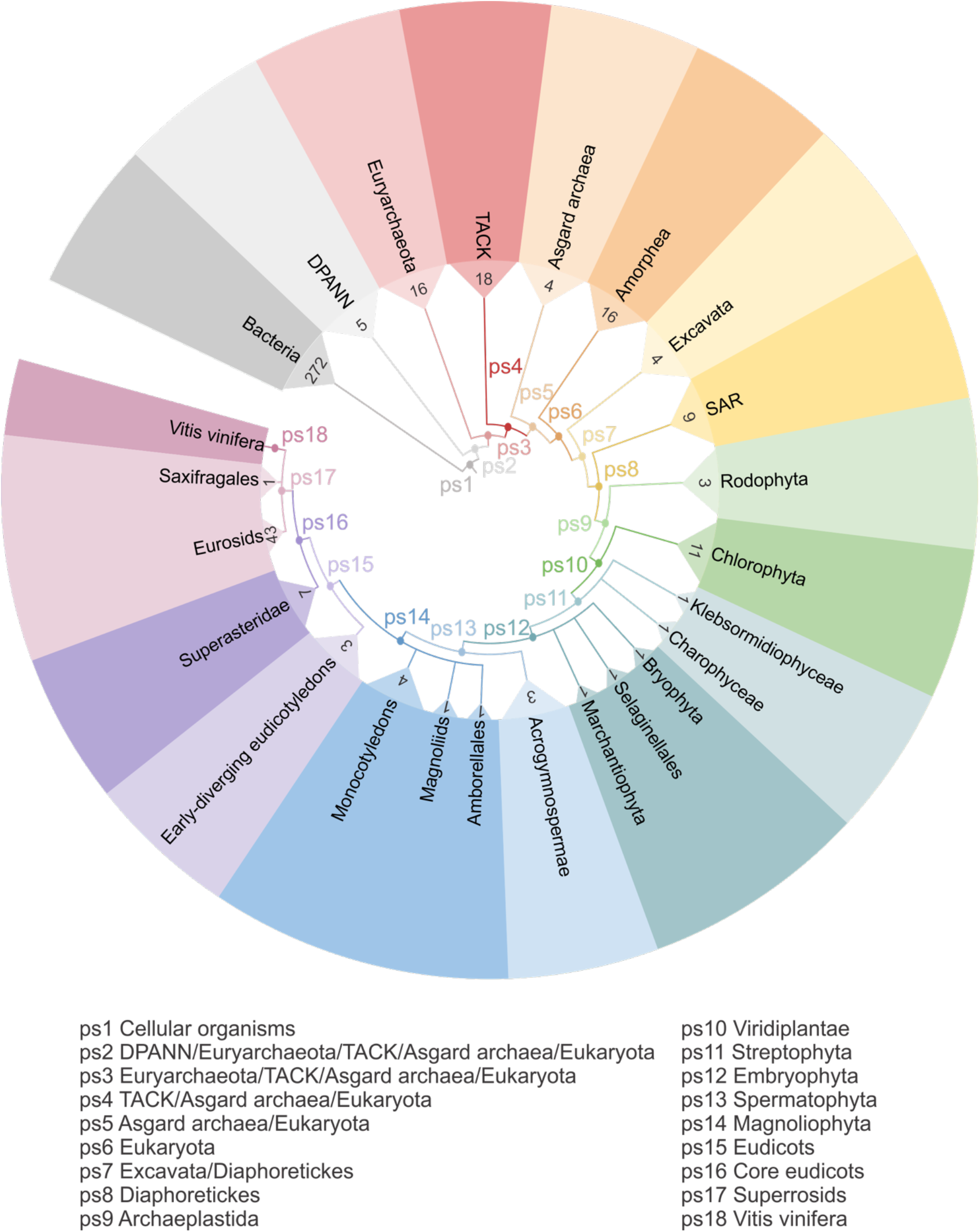
*Vitis vinifera* consensus phylogeny used for the phylostratigraphic analysis. The consensus phylogeny covers divergence from the last common ancestor of cellular organisms to *V. vinifera* as a focal organism. Phylogeny is constructed based on the relevant phylogenetic literature, importance of evolutionary transitions and availability of reference genomes. Eighteen internodes (phylostrata) are marked as ps1 – ps18. The numbers on the terminal nodes represent the number of species in the matching node and correspond to the genomes used to populate the reference database for sequence similarity searches.

**Supplementary Figure 4.**
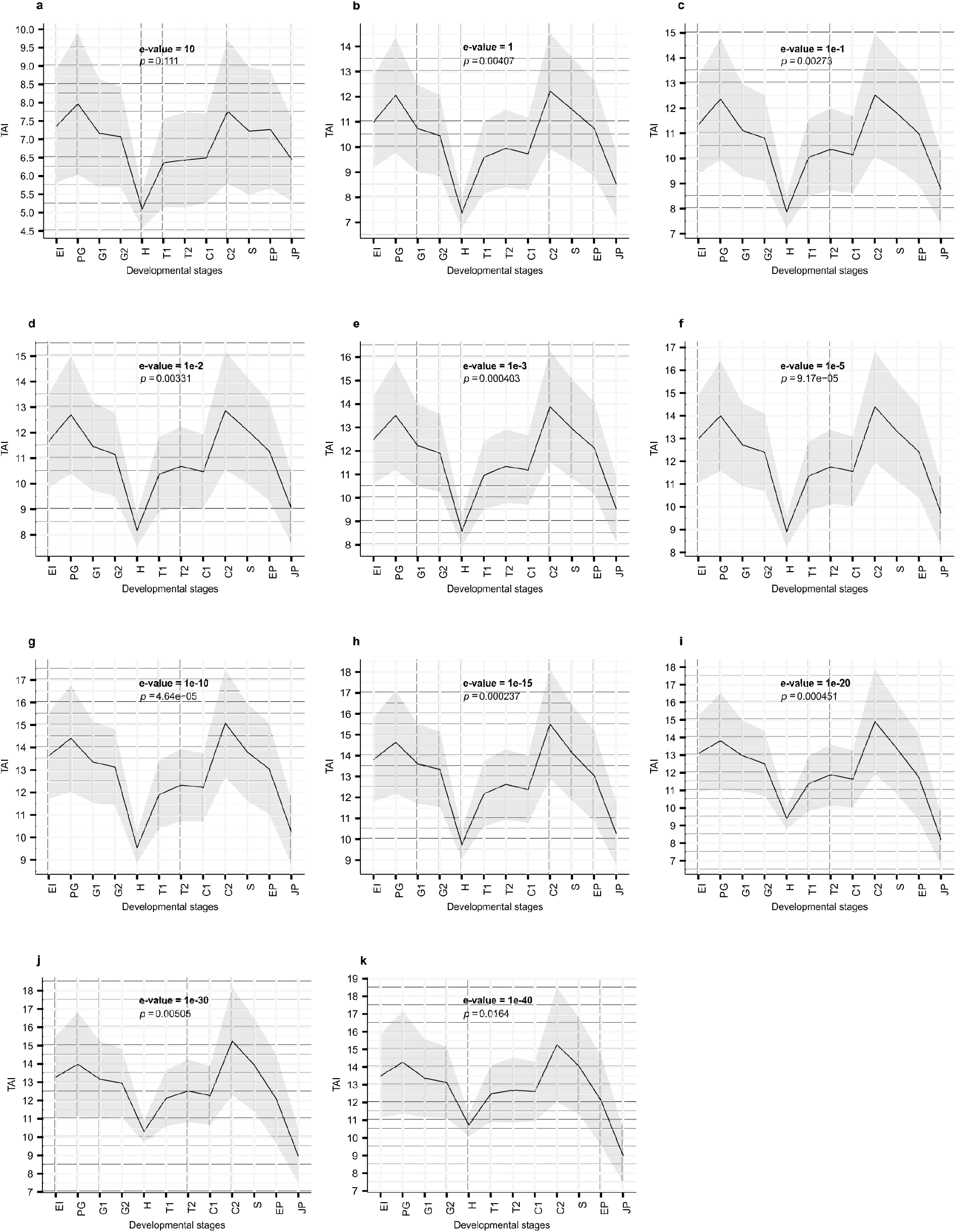
Transcriptome age index (TAI) profiles of *Vitis vinifera* somatic embryogenesis calculated from phylostratigraphy maps obtained with different blastp e-value thresholds. a, e-value = 10 (n = 29,659); **b**, e-value = 1 (n = 29,625); **c**, e-value = 10e-1 (n = 29,502); **d**, e-value = 10e-2 (n = 29,542); **e**, e-value = 10e-3 (n = 29,496); **f**, e-value = 10e-5 (n = 29,408); **g**, e-value = 10e-10 (n = 29,123); **h**, e-value = 10e-15 (n = 28,771); **i**, e-value = 10e-20 (n = 28,382); **j**, e-value = 10e-30 (n = 27,529); **k**, e-value = 10e-40 (n = 26,550). The *p* values shown were calculated using the flat line test while the grey shaded area represents ± one standard deviation estimated using permutation.

**Supplementary Figure 5.**
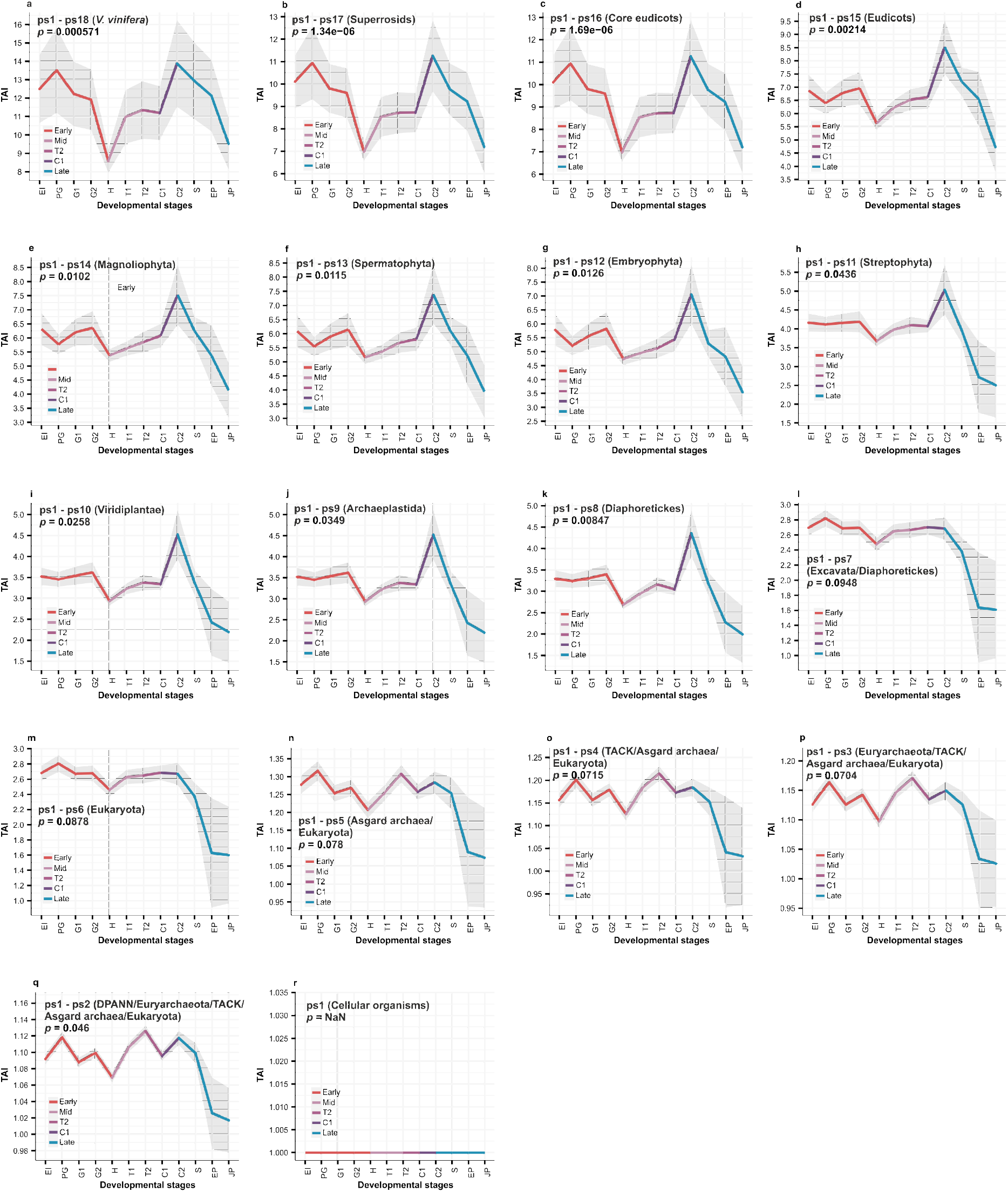
The hourglass pattern is significant from the origin of Diaphoretickes at ps8. Transcriptome age indices (TAI) were calculated using reduced datasets obtained by increasingly removing genes from the youngest phylostratum. This process was repeated until only genes from the oldest phylostratum (ps1) remained. **a**, ps1-ps18 (n=29,623); **b**, ps1-ps17 (n = 26,316); **c**, ps1-ps16 (n = 26,081); **d**, ps1-ps15 (n = 25,881); **e**, ps1-ps14 (n = 25,586); **f**, ps1-ps13 (n = 24,813); **g**, ps1-ps12 (n = 23,757); **h**, ps1-ps11 (n = 21,762); **i**, ps1-ps10 (n = 19,982); **j**, ps1-ps9 (n = 19,059); **k**, ps1-ps8 (n = 18,965); **l**, ps1-ps7 (n = 18,573); **m**, ps1-ps6 (n = 18,439); **n**, ps1-ps5 (n = 12,279); **o**, ps1-ps4 (n = 12,161); **p**, ps1-ps3 (n = 12,108); **q**, ps1-ps2 (n = 11,993); **r**, ps1 (n = 11,789). The *p* values were calculated using the flat line test while the grey shaded area represents ± one standard deviation estimated using permutation analysis (see Material and Methods). Expression phases of SE, as depicted in Fig. 1a, are color coded: early (red), mid, T2 and C1 (different shades of purple), and late (blue).

**Supplementary Figure 6.**
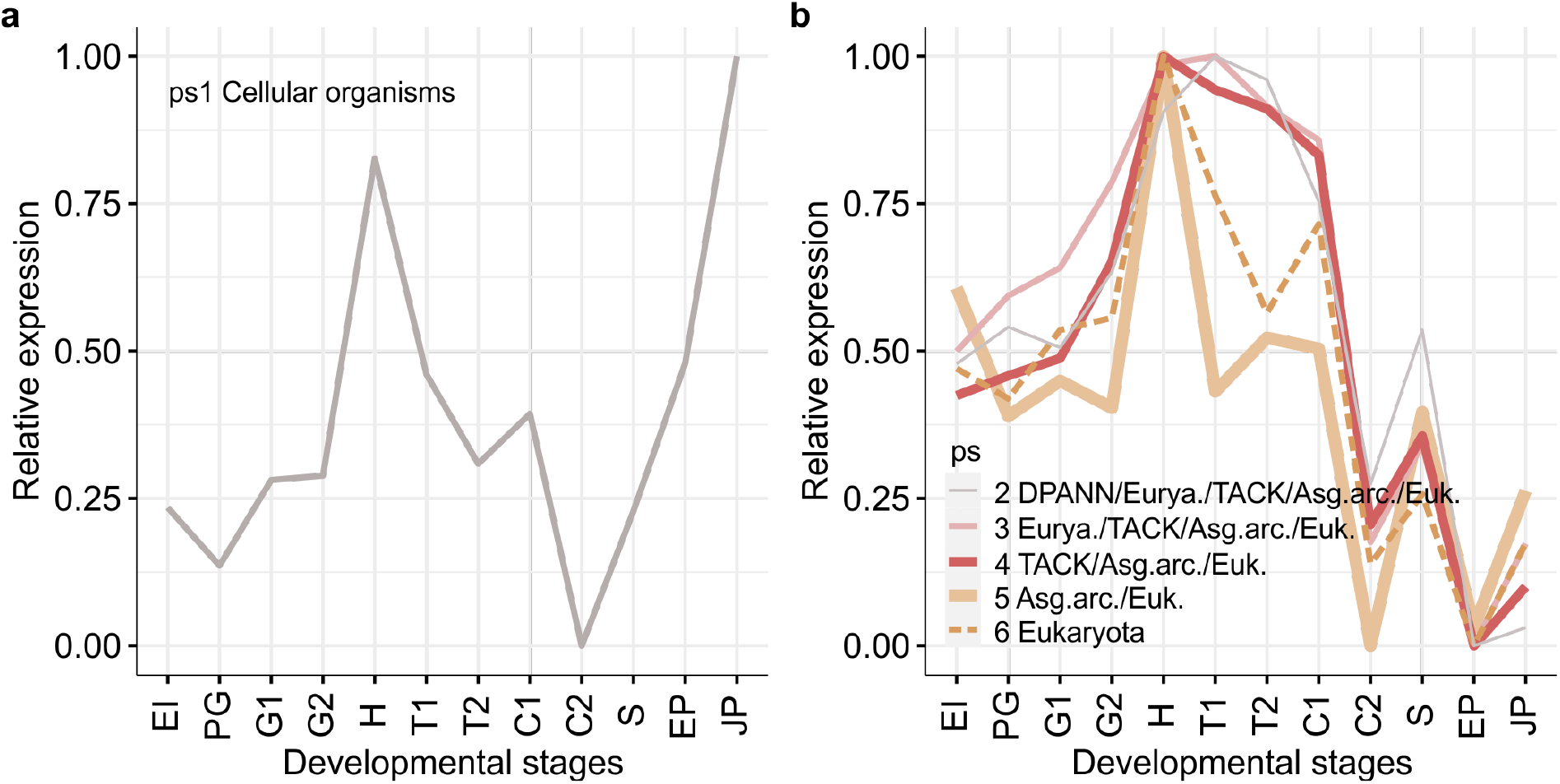
Relative expression (RE) profiles along *V. vinifera* somatic embryogenesis. Genes from phylostrata ps1 to ps6 are shown. **a**, RE profiles of the genes from Cellular organisms (ps1). **b**, RE profiles of the genes from DPANN (ps2) to Eukaryota (ps6). These older genes (ps1-ps6) generally peak at the heart stage.

**Supplementary Figure 7.**
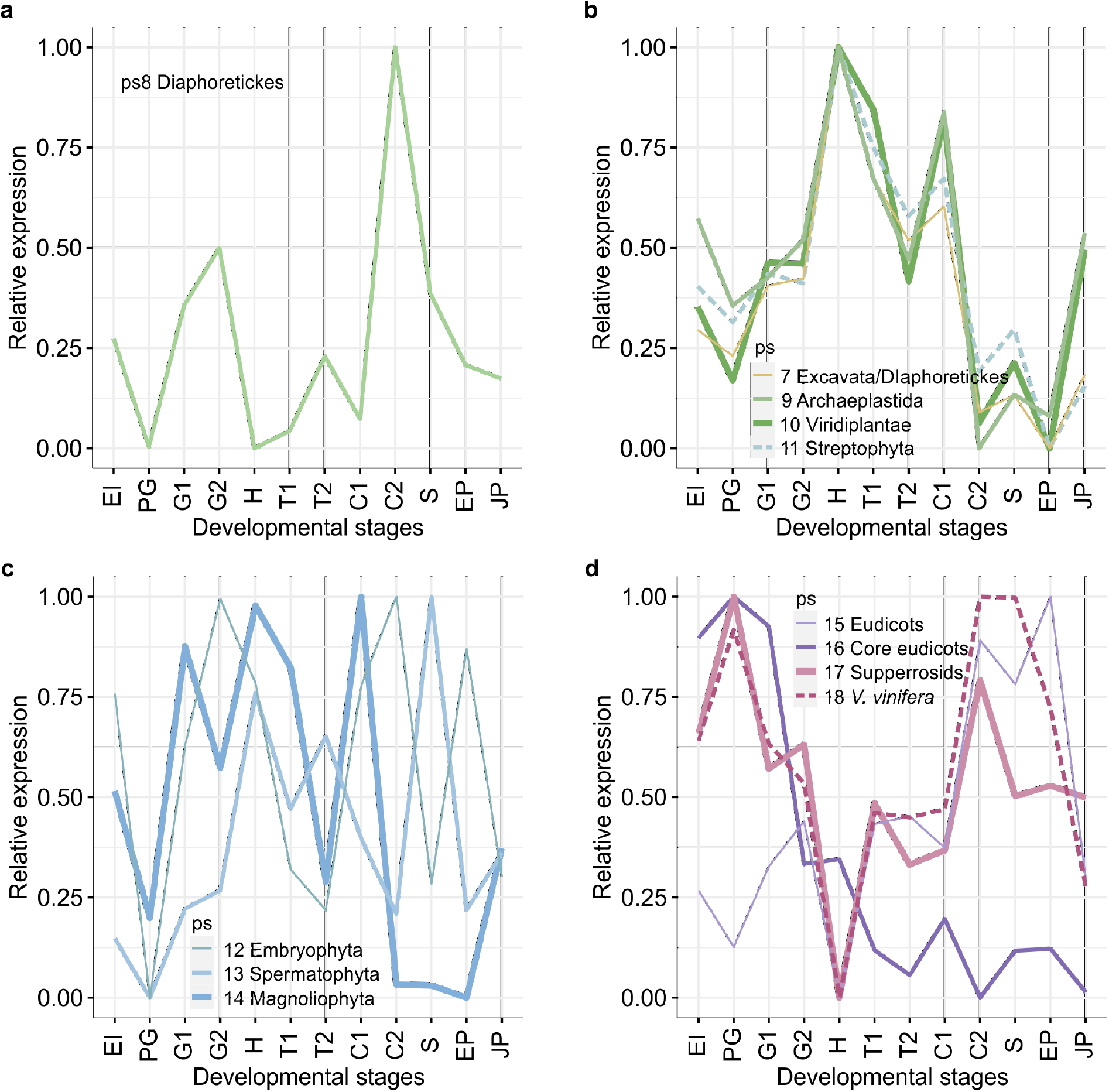
Relative expression (RE) profiles along *V. vinifera* somatic embryogenesis. Genes from phylostrata ps7 to ps18 are shown **a**, RE profile of the genes from Diaphoretickes (ps8). **b**, RE profiles of the genes from Excavata (ps7) and Archaeplastida (ps9) to Streptophyta (ps11). **c**, RE profiles of the genes from Embryophyta (ps12) to Magnoliophyta (ps14). **d**, RE profiles of the genes from Eudycots (ps15) to *V. vinifera* (ps18).

**Supplementary Figure 8.**
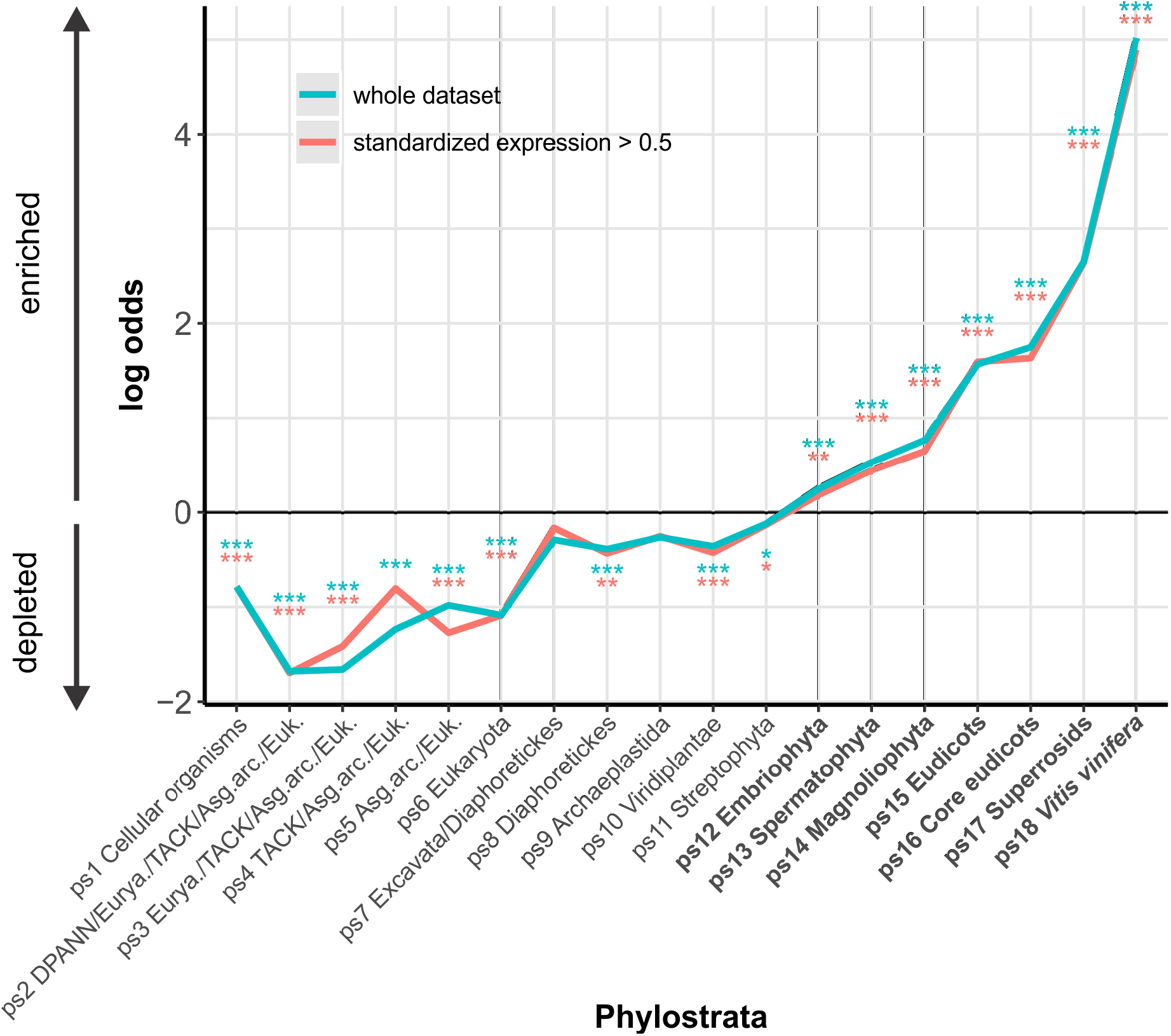
*V. vinifera* genes that emerged during the diversification of land plants tend to be functionally unstudied. The enrichment of genes with unknown function across phylostrata (n = 11,224, blue line), and for the subset of genes which standardized expression are at least in one developmental stage 0.5 or more above the median (n = 8,070, red line). We tested the enrichments, i.e., the significance of deviations from the expected values, by two-tailed hypergeometric test and *p* values are corrected for multiple comparisons at 0.05 level (*p < 0.05; **p < 0.01; ***p < 0.001). The enrichments are shown as log-odds (y-axis).

